# Unearthing the microbial ecology of soil carbon cycling with DNA-SIP

**DOI:** 10.1101/022483

**Authors:** Charles Pepe-Ranney, Ashley N Campbell, Chantal Koechli, Sean Berthrong, Daniel H Buckley

## Abstract

We explored the microbial contributions to decomposition using a sophisticated approach to DNA Stable Isotope Probing (SIP). Our experiment evaluated the dynamics and ecological characteristics of functionally defined microbial groups that metabolize labile and structural C in soils. We added to soil a complex amendment representing plant derived organic matter substituted with either ^13^C-xylose or ^13^C-cellulose to represent labile and structural C pools derived from abundant components of plant biomass. We found evidence for ^13^C-incorporation into DNA from ^13^C-xylose and ^13^C-cellulose in 49 and 63 operational taxonomic units (OTUs), respectively. The types of microorganisms that assimilated ^13^C in the ^13^C-xylose treatment changed over time being predominantly *Firmicutes* at day 1 followed by *Bacteroidetes* at day 3 and then *Actinobacteria* at day 7. These ^13^C-labeling dynamics suggest labile C traveled through different trophic levels. In contrast, microorganisms generally metabolized cellulose-C after 14 days and did not change to the same extent in phylogenetic composition over time. Microorganisms that metabolized cellulose-C belonged to poorly characterized but cosmopolitan soil lineages including *Verrucomicrobia*, *Chloroflexi* and *Planctomycetes.* We show that microbial life history traits are likely to constrain the diversity of microorganisms that participate in the soil C-cycle.

---

Abbreviations: C, Carbon; OTU, Operational Taxonomic Unit; SOM, Soil Organic Matter; BD, Buoyand Density; SIP, Stable Isotope Probing

## Introduction

Soils worldwide contain 2,300 Pg of carbon (C) which accounts for nearly 80% of the C present in the terrestrial biosphere [1, 2]. Soil microorganisms drive C flux through the terrestrial biosphere and C respiration by soil microorganisms produces annually tenfold more CO_2_ than fossil fuel emissions [3]. Despite the contribution of microorganisms to global C flux, many global C models ignore the diversity of microbial physiology [4–6] and we still know little about the ecophysiology of soil microorganisms. Characterizing the ecophysiology of microbes that mediate C decomposition in soil has proven difficult due to their overwhelming diversity. Such knowledge should assist the development and refinement of global C models [7–10].

The degradative succession hypothesis is a simple framework that explains the impact of microbial ecophysiology on the decomposition of plant biomass. Most plant C is comprised of cellulose (30–50%) followed by hemicellulose (20–40%), and lignin (15–25%) [11]. Hemicellulose, being the most soluble, degrades in the early stages of decomposition. Xylans are often an abundant component of hemicellulose, and xylose is often the most abundant sugar in hemicellulose, comprising as much as 60–90% of xylan in some plants (e.g switchgrass [12]). The degradative succession hypothesis posits that fast growing organisms proliferate in response to the labile fraction of plant biomass such as sugars [13, 14] followed by slow growing organisms that target structural C such as cellulose [13]. Evidence to support the degradative succession hypothesis comes from observing soil respiration dynamics and characterizing microorganisms cultured at different stages of decomposition. Microorganisms that consume labile C in the form of sugars proliferate during the initial stages of decomposition [15, 16], and metabolize as much as 75% of sugar C during the first 5 days [17]. In contrast, cellulose decomposition proceeds more slowly with rates increasing for approximately 15 days while degradation continues for 30–90 days [17, 18]. This hypothesis is generally consistent with the common categorization of soil microorganisms as either fast growing copiotrophs or slow growing oligotrophs [19]. The degree to which the degradative succession hypothesis presents an accurate model of litter decomposition has been questioned [20–22] and it’s clear that we need new: approaches to dissect microbial contributions to C transformations in soils.

Though microorganisms mediate 80–90% of the soil C-cycle [23, 24], and microbial community composition can account for significant variation: in C mineralization [25], terrestrial C-cycle models rarely consider the community composition of soils [26, 27]. Variation in microbial community composition can be linked effectively to rates of soil processes when diagnostic genes for specific functions: are available (e.g. nitrogen fixation [28]). However, the lack of diagnostic genes for describing soil-C transformations has limited progress in characterizing the contributions of individual microorganisms to decomposition. Remarkably, we still lack basic information on the physiology and ecology of the majority of organisms that live in soils. For example, contributions to soil processes remain uncharacterized for cosmopolitan bacterial phyla in soil such as *Acidobacteria, Chloroflexi, Planctomycetes,* and *Verrucomicrobia.* These phyla combined can comprise 32% of soil microbial communities (based on surveys of the SSU rRNA genes in soil) [29, 30].

Nucleic acid stable-isotope probing (SIP) links genetic identity and activity without the need diagnostic genetic markers or cultivation and has expanded our knowledge of microbial processes [31]. Nucleic acid SIP has notable complications, however, including the need to add large amounts of labeled substrate [32], label dilution resulting in partial labeling of nucleic acids [32], the potential for cross-feeding and secondary label incorporation [33], and variation in genome G+C content [34]. As a result, most applications of SIP have targeted specialized microorganisms (for instance, methylotrophs [35], syntrophs [36], or microorganisms that target pollutants [37]). Exploring the soil-C cycle with SIP has proven to be more challenging because SIP has lacked the resolution necessary to characterize the specific contributions of individual microbial groups to the decomposition of plant biomass. High throughput DNA sequencing technology, however, improves the resolving: power of SIP [38]. It is now possible to use far less isotopically labeled substrate resulting in more environmentally realistic experimental conditions. It is also possible to sequence rRNA genes from numerous density gradient fractions across multiple samples thereby increasing the resolution of a typical nucleic acid SIP experiment [39]. We have employed such a high resolution DNA stable isotope probing approach to explore the assimilation of both xylose and cellulose into bacterial DNA in an agricultural soil.

We added to soil a complex amendment representative of organic matter derived from fresh plant biomass. All treatments received the same amendment but the identity of isotopically labeled substrates was varied between treatments. Specifically, we set up a control treatment where all components were unlabeled, a treatment with ^13^C-xylose instead of unlabeled xylose, and a treatment with ^13^C-cellulose instead of unlabeled cellulose. Soil was sampled at days 1, 3, 7, 14, and 30 and we identified microorganisms that assimilated ^13^ C into DNA at each point in time. We designed the experiment to test of the degradative succession hypothesis as it applies to soil bacteria, to identify soil bacteria that metabolize xylose and cellulose, and to characterize temporal dynamics of xylose and cellulose metabolism in soil.

## Results

After adding the organic matter amendment to soil, we tracked the flow of ^13^C from ^13^C-xylose or ^13^C-cellulose into microbial DNA over time using DNA-SIP (Figure S1). The amendment consisted of compounds representative of plant biomass including cellulose, lignin, sugars found in hemicellulose, amino acids, and inorganic nutrients (see Supplemental Information (SI)). The amendment was added at 2.9 mg C g^−1^ soil dry weight (d.w.), and this comprised 19% of the total C in the soil. The cellulose-C (0.88 mg C g^−1^ soil d.w.) and xyloseC (0.42 mg C g^−1^ soil d.w.) in the amendment comprised 6% and 3% of the total C in the soil, respectively. The soil microbial community respired 65% of the xylose within one day and 29% of the added xylose remained in the soil at day 30 (Figure S2). In contrast, cellulose-C declined at a rate of approximately 18 *μ*g C d ^−1^ g^−1^ soil d.w. and 40% of added cellulose-C remained in the soil at day 30 (Figure S2).

**Community-level signal of ^13^C-assimilation in relation to substrate and time.** We assessed assimilation of ^13^C into microbial DNA by comparing the SSU rRNA gene sequence composition of SIP density gradient fractions between ^13^ C treatments and the unlabeled control (see Methods and SI). Our main focus is to identify evidence of isotope incorporation into the DNA of specific OTUs (as described below), but it is instructive to begin by observing overall patterns of variance in the SSU rRNA gene sequence composition of gradient fractions. In the unlabeled control treatment, fraction density represented the majority of the variance in SSU rRNA gene composition (Figure 1). This result is expected because Genome G+C content correlates positively with DNA buoyant density and influences SSU rRNA gene composition in gradient fractions [34]. In contrast, isotope assimilation into DNA will cause variation in gene sequence composition between corresponding density fractions from controls and labeled treatments. For example, the SSU rRNA gene composition in gradient fractions from the ^13^C-cellulose treatment deviated from corresponding control fractions on days 14 and 30 and this difference was observed only in the high density fractions (>1.7125 g mL^−1^, Figure 1). Likewise, SSU rRNA gene composition in gradient fractions from the ^13^C-xylose treatment also deviated from corresponding control fractions but on days 1, 3, and 7 as opposed to 14 and 30 (Figure 1). The ^13^C-cellulose and ^13^C-xylose treatments also differed from each other in corresponding high density gradient fractions indicating that different microorganisms were labeled across time these treatments (Figure 1). These results are generally consistent with predictions of the degradative succession hypothesis.

**Fig. 1.**
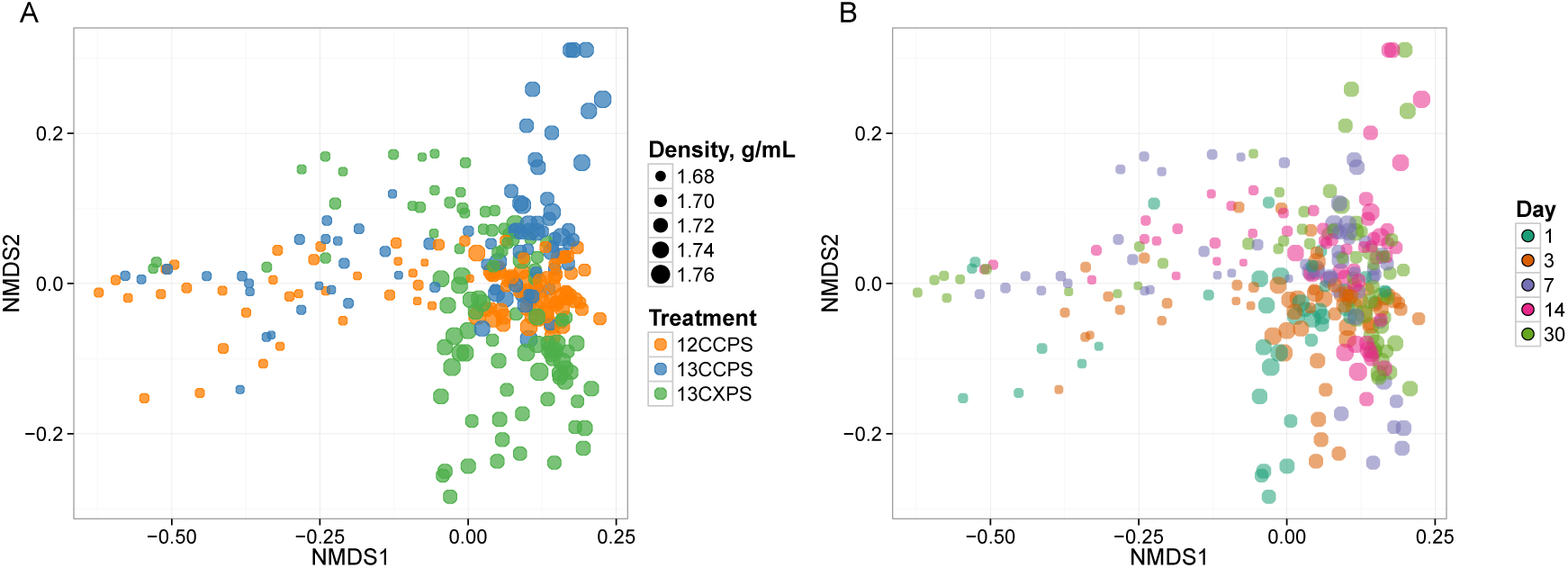
NMDS ordination of SSU rRNA gene sequence composition in gradient fractions shows that variation between fractions is correlated with fraction density, isotopic labeling, and time. Dissimilarity in SSU rRNA gene sequence composition was quantified using the weighted UniFrac metric. SSU rRNA gene sequences were surveyed in twenty gradient fractions at each sampling point for each treatment (Figure S1). ^13^C-labeling of DNA is apparent because the SSU rRNA gene sequence composition of gradient fractions from ^13^C and control treatments differ at high density. Each point on the NMDS plot represents one gradient fraction. SSU rRNA gene sequence composition differences between gradient fractions were quantified by the weighted Unifrac metric. The size of each point is positively correlated with density and colors indicate the treatment (A) or day (B).

We can observe further differences in the pattern of isotope incorporation over time for each treatment. For example the SSU rRNA gene sequence composition in the ^13^C-cellulose treatment was similar on days 14 and 30 in corresponding high density fractions indicating similar patterns of isotope incorporation into DNA on the days. In contrast, in the ^13^C-xylose treatment, the SSU rRNA gene composition varied between days 1, 3, and 7 in corresponding high density fractions indicating different pattenrs of isotope incorporation into DNA on these days. In the ^13^C-xylose treatment on days 14 and 40 the SSU gene composition was similar to control on days 14 and 30 for corresponding high density fractions (Figure 1) indicating that ^13^C was no longer detectable in bacterial DNA on these days for this treatment. These results show that the dynamics of isotope incorporation into DNA varied considerably for organisms that assimilated C from either xylose or cellulose.

**Temporal dynamics of OTU relative abundance in non-fractionated DNA from soil.** We monitored the soil microbial community over the course of the experiment by surveying SSU rRNA genes in non-fractionated DNA from the soil. The SSU rRNA gene composition of the non-fractionated DNA changed with time (Figure S3, P-value = 0.023, R^2^ = 0.63, Adonis test [40]). In contrast, the microbial community could not be shown to change with treatment (P-value 0.23, Adonis test) (Figure S3). The latter result demonstrates the substitution of ^13^C-labeled substrates for unlabeled equivalents could not be shown to alter the soil microbial community composition. Twenty-nine OTUs exhibited sufficient statistical evidence (adjusted P-value <0.10, Wald test) to conclude they changed in relative abundance in the non-fractionated DNA over the course of the experiment (Figure S4). When SSU rRNA gene abundances were combined at the taxonomic rank of “class”, the classes that changed in abundance (adjusted P-value < 0.10, Wald test) were the *Bacilli* (decreased), *Flavobacteria* (decreased), *Gammaproteobacteria* (decreased), and *Herpetosiphonales* (increased) (Figure S5). Of the 29 OTUs that changed in relative abundance over time, 14 putatively incorporated ^13^C into DNA (see below and Figure S4). OTUs that likely assimilated ^13^C from ^13^C-cellulose tended to increase in relative abundance with time whereas OTUs that assimilated ^13^C from ^13^C-xylose tended to decrease (Figure S6). OTUs that responded to both substrates did not exhibit a consistent relative abundance response over time as a group (Figure S4 and S6).

**Changes in the phylogenetic composition of ^13^C-labeled OTUs with substrate and time.** If an OTU exhibited strong evidence for assimilating ^13^C into DNA, we refer to that OTU as a “responder” (see Methods and SI for our operational definition of “responder”). The SSU rRNA gene sequences produced in this study were binned into 5,940 OTUs and we assessed evidence of ^13^C-labeling from both ^13^C-cellulose and ^13^C-xylose for each OTU. Forty-one OTUs responded to ^13^C-xylose, 55 OTUs responded to ^13^C-cellulose, and 8 OTUs responded to both xylose and cellulose (Figure 2, Figure 3, Figure S7, Table S1, and Table S2). The number of xylose responders peaked at days 1 and 3 and declined with time. In contrast, the number of cellulose responders increased with time peaking at days 14 and 30 (Figure S8).

**Fig. 2.**
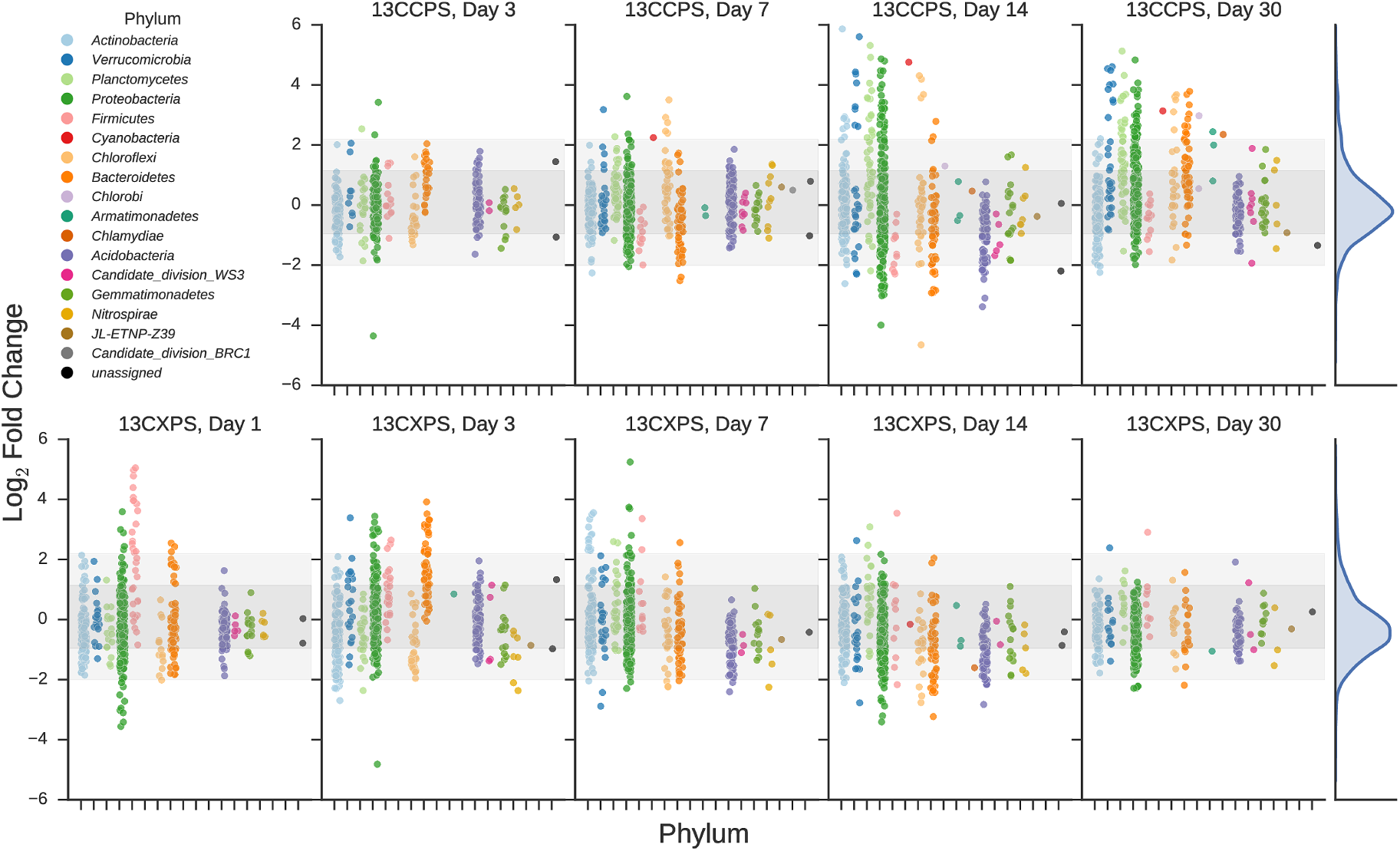
Enrichment of OTUs in either ^13^C-cellulose (13CCPS, upper panels) or ^13^C-xylose (13CXPS, bottom panels) treatments relative to control, expressed as LFC (see Methods). Each point indicates the LFC for a single OTU. High enrichment values indicate an OTU is likely ^13^C-labeled. Different colors represent different phyla and different panels represent different days. The final column shows the frequency distribution of LFC values in each row. Within each panel, shaded areas are used to indicate one standard deviation (dark shading) or two standard deviations (light shading) about the mean of all LFC values.

**Fig. 3.**
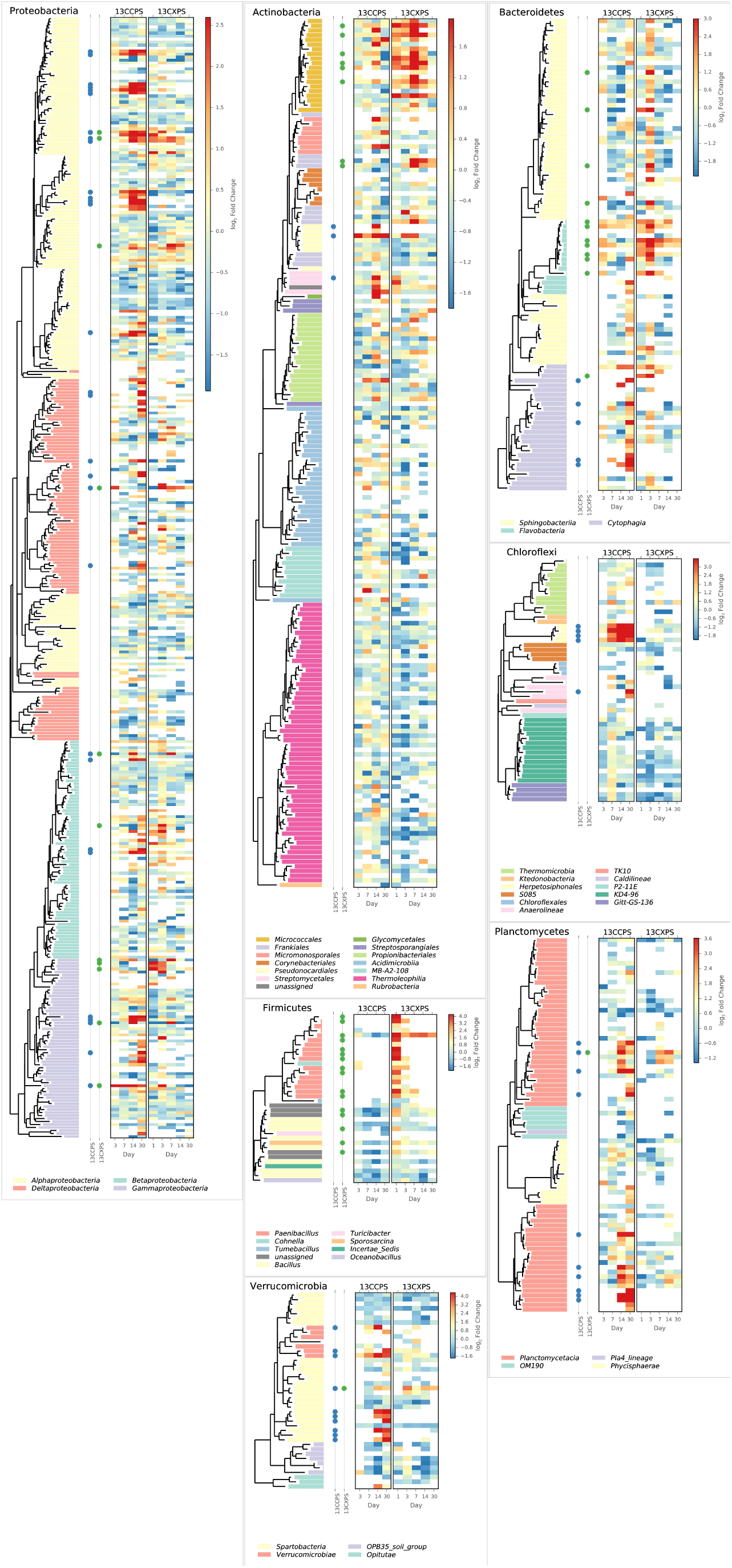
Phylogenetic position of cellulose responders and xylose responders in the context of all OTUs that passed sparsity independent filtering criteria (see Methods). Only those phyla that contain responders are shown. Colored dots are used to identify xylose responders (green) and cellulose responders (blue). The heatmaps indicate enrichment in high density fractions relative to control (represented as LFC) for each OTU in response to both ^13^C-cellulose (13CCPS, leftmost heatmap) and ^13^C-xylose (13CXPS, rightmost heatmap) with values for different days in each heatmap column. High enrichment values (represented as LFC) provide evidence of ^13^C-labeled DNA.

The phylogenetic composition of xylose responders changed with time (Figure 2 and Figure 4) and 86% of xylose responders shared > 97% SSU rRNA gene sequence identity with bacteria cultured in isolation (Table S1). On day 1, *Bacilli* OTUs represented 84% of xylose responders (Figure 4) and the majority of these OTUs were closely related to cultured representatives of the genus *Paenibacillus* (Table S1, Figure 3). For example, “OTU.57” (Table S1), annotated as *Paenibacillus*, had a strong signal of ^13^C-labeling at day 1 coinciding with its maximum relative abundance in non-fractionated DNA. The relative abundance of “OTU.57” declined until day 14 and “OTU.57” did not appear to be ^13^C-labeled after day 1 (Figure S9). On day 3, *Bacteroidetes* OTUs comprised 63% of xylose responders (Figure 4) and these OTUs were closely related to cultured representatives of the *Flavobacteriales* and *Sphingobacteriales* (Table S1, Figure 3). For example, “OTU.14”, annotated as a flavobacterium, had a strong signal for ^13^C-labeling in the ^13^C-xylose treatment at days 1 and 3 coinciding with its maximum relative abundance in non-fractionated DNA. The relative abundance of “OTU.14” then declined until day 14 and did not show evidence of ^13^C-labeling beyond day 3 (Figure S9). Finally, on day 7, *Actinobacteria* OTUs represented 53% of the xylose responders (Figure 4) and these OTUs were closely related to cultured representatives of *Micrococcales* (Table S1, Figure 3). For example, “OTU.4”, annotated as *Agromyces,* had signal for ^13^C-labeling in the ^13^C-xylose treatment on days 1, 3 and 7 with the strongest evidence of ^13^C-labeling at day 7 and did not appear ^13^C-labeled at days 14 and 30. The relative abundance of “OTU.4” in non-fractionated DNA increased until day 3 and then declined until day 30 (Figure S9). *Proteobacteria* were also common among xylose responders at day 7 where they comprised 40% of xylose responder OTUs. Notably, *Proteobacteria* represented the majority (6 of 8) of OTUs that responded to both cellulose and xylose (Figure S7).

**Fig. 4.**
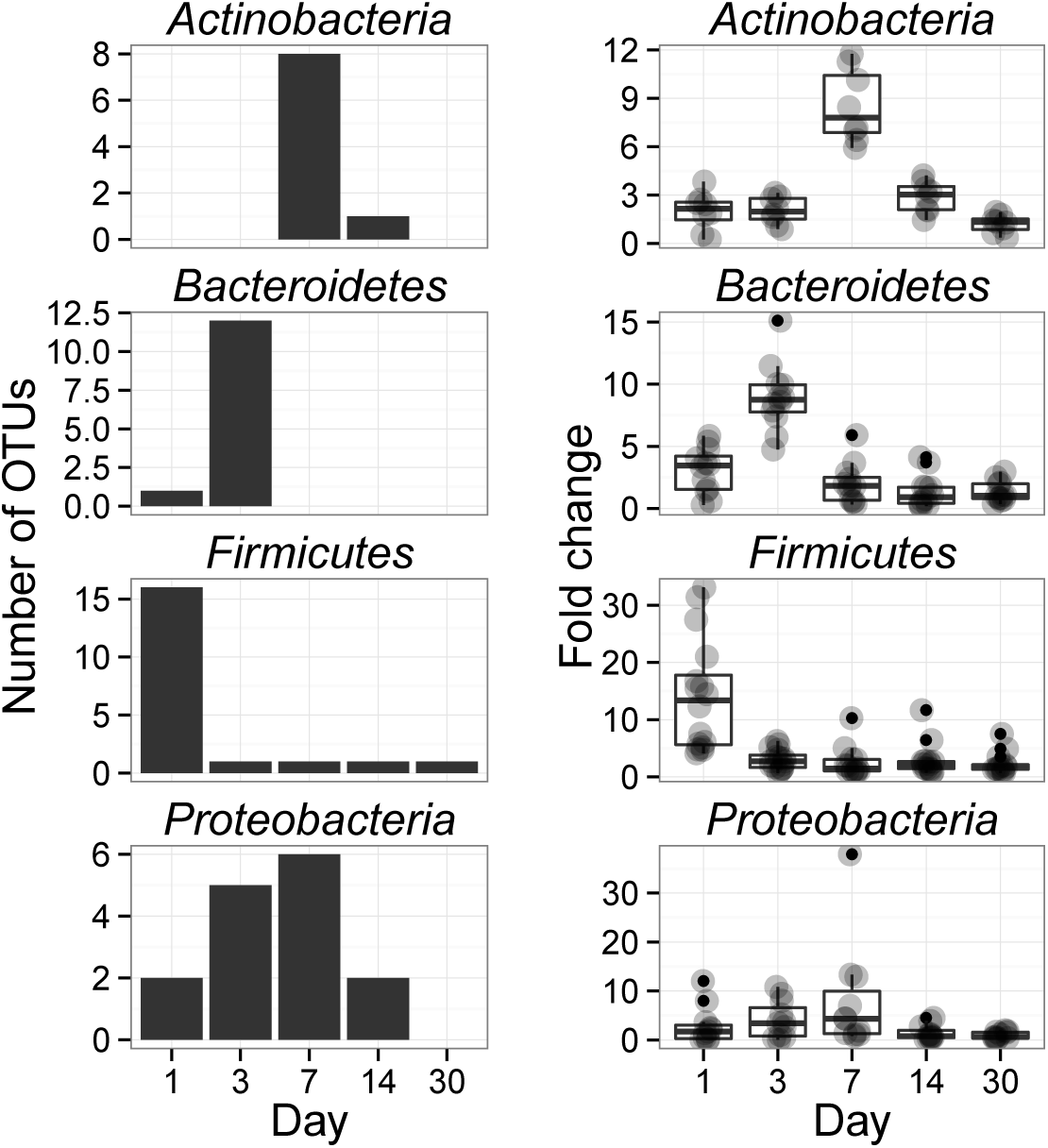
Xylose reponders in the *Actinobacteria, Bacteroidetes, Firmicutes* exhibit distinct temporal dynamics of ^13^C-labeling. The left column shows counts of ^13^C-xylose responders in the *Actinobacteria, Bacteroidetes, Firmicutes* and *Proteobacteria* at days 1, 3, 7 and 30. The right panel shows OTU enrichment in high density gradient fractions (gray points, expressed as fold change) for responders as well as a boxplot for the distribution of fold change values (The box extends one interquartile range, whiskers extend 1.5 times the IR, and small dots are outliers (i.e. beyond 1.5 times the IR)). Each day in the right column shows all responders (i.e. OTUs that responded to xylose at any point in time). High enrichment values indicates OTU DNA is likely ^13^C-labeled.

The phylogenetic composition of cellulose responders did not change with time to the same extent as the xylose responders. Also, in contrast to xylose responders, cellulose responders often were not closely related (< 97% SSU rRNA gene sequence identity) to cultured isolates. Both the relative abundance and the number of cellulose responders increased over time peaking at days 14 and 30 (Figure 2, Figure S8, and Figure S6). Cellulose responders belonged to the *Proteobacteria* (46%), *Verrucomicrobia* (16%), *Planctomycetes* (16%), *Chloroflexi* (8%), *Bacteroidetes* (8%), *Actinobacteria* (3%), and *Melainabacteria* (1 OTU) (Table S2).

The majority (85%) of cellulose responders outside of the *Proteobacteria* shared < 97% SSU rRNA gene sequence identity to bacteria cultured in isolation. For example, 70% of the *Verrucomicrobia* cellulose responders fell within unidentified *Spartobacteria* clades (Figure 3), and these shared < 85% SSU rRNA gene sequence identity to any characterized isolate. The *Spartobacteria* OTU “OTU.2192” exemplified many cellulose responders (Table S2, Figure S9). “OTU.2192” increased in non-fractionated DNA relative abundance with time and evidence for ^13^C-labeling of “OTU.2192” in the ^13^C-cellulose treatment increased over time with the strongest evidence at days 14 and 30 (Figure S9). Most *Chloroflexi* cellulose responders belonged to an unidentified clade within the *Herpetosiphonales* (Figure 3) and they shared < 89% SSU rRNA gene sequence identity to any characterized isolate. Characteristic of *Chloroflexi* cellulose responders, “OTU.64” increased in relative abundance over 30 days and evidence for ^13^C-labeling of “OTU.64” in the ^13^C-cellulose treatment peaked days 14 and 30 (Figure S9). *Bacteroidetes* cellulose responders fell within the *Cytophagales* in contrast with *Bacteroidetes* xylose responders that belonged instead to the *Flavobacteriales* or *Sphingobacteriales* (Figure 3). *Bacteroidetes* cellulose responders included one OTU that shared 100% SSU rRNA gene sequence identity to a *Sporocytophaga* species, a genus known to include cellulose degraders. The majority (86%) of cellulose responders in the *Proteobacteria* were closely related (> 97% identity) to bacteria cultured in isolation, including representatives of the genera: *Cellvibrio, Devosia, Rhizobium,* and *Sorangium*, which are all known for their ability to degrade cellulose (Table S2). Proteobacterial cellulose responders belonged to *Alpha* (13 OTUs), *Beta* (4 OTUs), *Gamma* (5 OTUs), and *Deltaproteobacteria* (6 OTUs).

### Characteristics of cellulose and xylose responders

Cellulose responders, relative to xylose responders, tended to have lower relative abundance in non-fractionated DNA, demonstrated signal consistent with higher atom % ^13^C in labeled DNA, and had lower estimated *rrn* copy number (Figure 5). OTUs that assimilated C from either cellulose or xylose were also clustered phylogenetically (see below) indicating that these traits were not dispersed randomly across bacterial species.

**Fig. 5.**
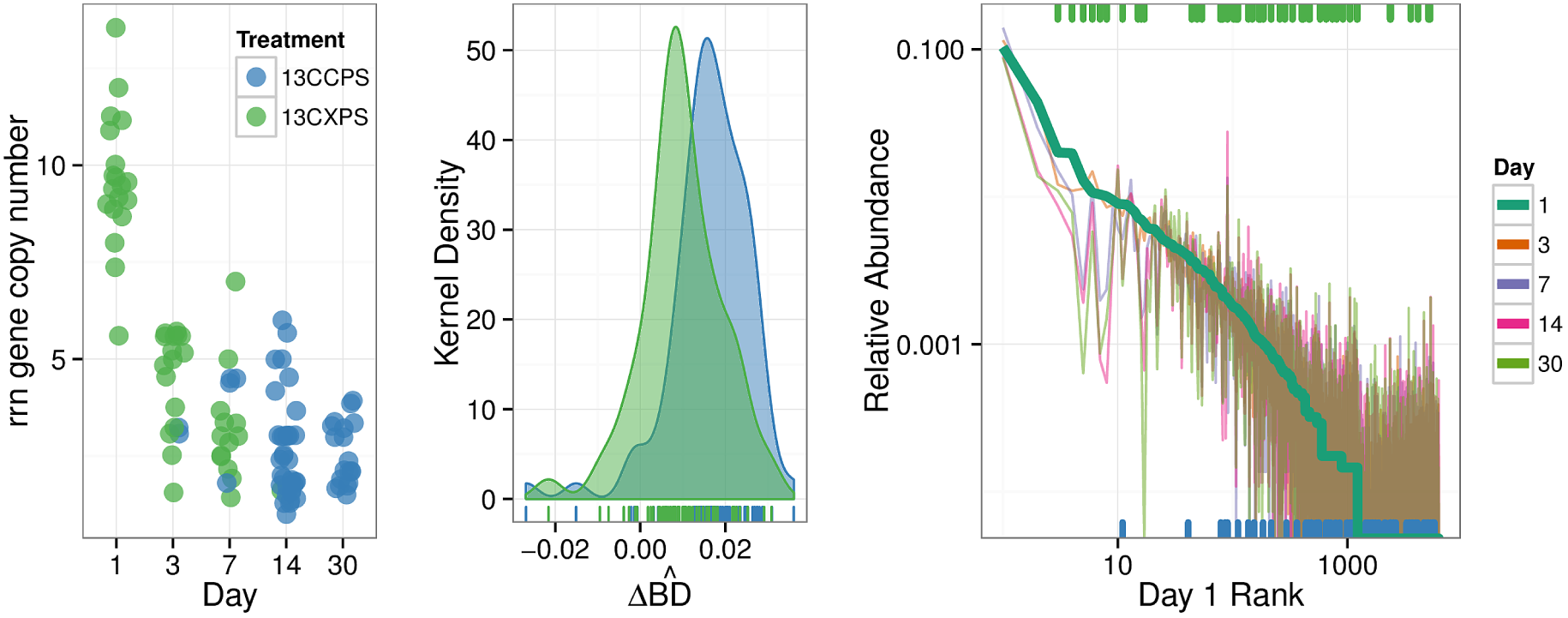
Characteristics of xylose responders (green) and cellulose responders (blue) based on estimated *rrn* copy number (A), 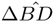 (B), and relative abundance in non-fractionated DNA (C). The estimated *rrn* copy number of all responders is shown versus time (A). Kernel density histogram of 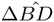 values shows cellulose responders had higher average 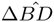 than xylose responders indicating higher average atom % ^13^C in OTU DNA (B). The final panel indicates the rank relative abundance of all OTUs observed in the non-fractionated DNA (C) where rank was determined at day 1 (bold line) and relative abundance for each OTU is indicated for all days by colored lines (see legend). Xylose responders (green ticks) have higher relative abundance in non-fractionated DNA than cellulose responders (green ticks). All ticks are based on day 1 relative abundance.

In the non-fractionated DNA, cellulose responders had lower relative abundance (1.2 × 10^−3^ (s.d. 3.8 × 10^−3^)) than xylose responders (3.5 × 10^−3^ (s.d. 5.2 × 10^−3^)) (Figure 4, P-value = 1.12 × 10^−5^, Wilcoxon Rank Sum test). Six of the ten most common OTUs observed in the non-fractionated DNA responded to xylose, and, seven of the ten most abundant responders to xylose or cellulose in the non-fractionated DNA were xylose responders.

DNA buoyant density (BD) increases in proportion to atom % ^13^C. Hence, the extent of ^13^C incorporation into DNA can be evaluated by the difference in BD between ^13^C-labeled and unlabeled DNA. We calculated for each OTU its mean BD weighted by relative abundance to determine its “center of mass” within a given density gradient. We then quantified for each OTU the difference in center of mass between control gradients and gradients from ^13^C-xylose or ^13^C-cellulose treatments (see SI for the detailed calculation, Figure S11). We refer to the change in center of mass position for an OTU in response to ^13^C-labeling as 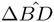. This value can be used to compare relative differences in ^13^C-labeling between OTUs. 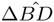values, however, are not comparable to the BD changes observed for DNA from pure cultures both because they are based on relative abundance in density gradient fractions (and not DNA concentration) and because isolated strains grown in uniform conditions generate uniformly labeled molecules while OTUs composed of heterogeneous strains in complex environmental samples do not. Cellulose responder 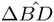 (0.0163 g mL^−1^ (s.d. 0.0094)) was greater than that of xylose responders (0.0097 g mL^−1^ (s.d. 0.0094)) (Figure 5, P-value = 1.8610 × 10^−6^, Wilcoxon Rank Sum test).

We predicted the *rrn* gene copy number for responders as described [41]. The ability to proliferate after rapid nutrient influx correlates positively to a microorganism’s *rrn* copy number [42]. Cellulose responders possessed fewer estimated *rrn* copy numbers (2.7 (1.2 s.d.)) than xylose responders (6.2 (3.4 s.d.)) ( P = 1.878 × 10^−9^, Wilcoxon Rank Sum test, Figure 5 and Figure S10). Furthermore, the estimated *rrn* gene copy number for xylose responders was inversely related to the day of first response (P = 2.02 × 10^−15^, Wilcoxon Rank Sum test, Figure S10, Figure 5).

We assessed phylogenetic clustering of ^13^ C-responsive OTUs with the Nearest Taxon Index (NTI) and the Net Relatedness Index (NRI) [43]. We also quantified the average clade depth of cellulose and xylose responders with the consenTRAIT metric [44]. Briefly, the NRI and NTI evaluate phylogenetic clustering against a null model for the distribution of a trait in a phylogeny. The NRI and NTI values are z-scores or standard deviations from the mean and thus the greater the magnitude of the NRI/NTI, the stronger the evidence for clustering (positive values) or overdispersion (negative values). NRI assesses overall clustering whereas the NTI assesses terminal clustering [45]. The consenTRAIT metric is a measure of the average clade depth for a trait in a phylogenetic tree. NRI values indicate that cellulose responders clustered overall and at the tips of the phylogeny (NRI: 4.49, NTI: 1.43) while xylose responders clustered terminally (NRI: -1.33, NTI: 2.69). The consenTRAIT clade depth for xylose and cellulose responders was 0.012 and 0.028 SSU rRNA gene sequence dissimilarity, respectively. As reference, the average clade depth as inferred from genomic analyses or growth in culture is approximately 0.017, 0.013 and 0.034 SSU rRNA gene sequence dissimilarity for arabinase (arabinose like xylose is a five C sugar found in hemicellulose), glucosidase and cellulase, respectively [44, 46]. These results indicate xylose responders form terminal clusters dispersed throughout the phylogeny while cellulose responders form deep clades of terminally clustered OTUs.

## Discussion

We highlight two key results with implications for understanding structure-function relationships in soils, and for applying DNA-SIP in future studies of the soil-C cycle. First, cellulose responders were members of physiologically undescribed taxonomic groups with few exceptions. This suggests that we have much to learn about the diversity of structural-C decomposers in soil before we can begin to assess how they are affected by climate change and land management. Second, the response to xylose was characterized by a succession in activity from *Paenibacillus* OTUs (day 1) to *Bacteroidetes* (day 3) and finally *Micrococcales* (day 7). This activity succession was mirrored by relative abundance profiles and may mark trophic-C exchange between these groups. While trophic exchange has been observed previously in DNA-SIP studies [35] most applications of DNA-SIP focus on proximal use of labeled substrates. However, with increased sensitivity, DNA-SIP is well suited to tracking C flows throughout microbial communities over time and is not limited only to observing the entry point for a given substrate into the soil C-cycle. Trophic interactions will critically influence how the global soil-C reservoir will respond to climate change [47] but we know little of biological interactions among soil bacteria. Often bacteria are cast as a single trophic level [48] but it may be appropriate to investigate the soil food web at greater granularity. Additionally, our results show that DNA-SIP results can change dramatically over time suggesting that multiple time points are necessary to rigorously and comprehensively describe which microorganisms consume ^13^C-labeled substrates in nucleic acid SIP incubations.

Microorganisms that consumed ^13^C-cellulose were seldom related closely to any physiologically characterized cultured isolates but were members of cosmopolitan phylogenetic groups in soil including *Spartobacteria*, *Planctomycetes*, and *Chloroflexi.* Often cellulose responders were less than 90% related to their closest cultured relatives showing that we can infer little, if anything at all, of their physiology from culture-based studies. Notably, many *Spartobacteria* were among the cellulose responder OTUs. This is particularly interesting as *Spartobacteria* are globally distributed and found in a variety of soil types [49]. These lineages may play important roles in global cellulose turnover (please see SI note 1 for further discussion of the phylogenetic affiliation of cellulose responders).

In addition to taxonomic identity, we quantified four ecological properties of microorganisms that were actively engaged in labile and structural C decomposition in our experiment: (1) time of activity, (2) estimated *rrn* gene copy number, (3) phylogenetic clustering, and (4) density shift in response to ^13^C-labeling. Labile C was consumed before structural C and these substrates were consumed by different microorganisms (Figure 1). This was expected and is consistent with the degradative succession hypothesis. Consumers of labile C had higher estimated *rrn* gene copy number than structural C consumers (Figure 5). *rrn* copy number is positively correlated with the ability to resuscitate quickly in response to nutrient influx [42] which may be the advantage that enabled xylose responders to rapidly consume xylose. Both xylose and cellulose responders were terminally clustered phylogenetically suggesting that the ability to use these substrates was phylogenetically constrained. Although labile C consumption is generally considered to be mediated by a diverse set of microorganisms, we found that xylose responders at day 1 were mainly members of one genus, *Paenibacillus*. Our results suggests that life-history traits such as the ability to resuscitate quickly and/or grow rapidly may be more important in determining the diversity of microorganisms that actually mediate a given process than the genomic potential for substrate utilization (see SI note 2 for further discussion with respect to soil-C modelling). And last, labile C consumers, in contrast to structural C consumers, had lower 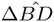 in response to ^13^C-labeling. This result suggests that labile C consumers were generalists, assimilating C from a variety of sources both labeled and unlabeled, while structural C consumers were more likely to be specialists and more closely associated with C from a single source.

We propose that the temporal fluctuations in ^13^C-labeling in the ^13^C-xylose treatment are due to trophic exchange of ^13^C. Alternatively, the temporal dynamics could be caused by microorganisms tuned to different substrate concentrations and/or cross-feeding. However, trophic exchange would explain well the precipitous drop in abundance of *Paenibacillus* after day 1 with subsequent ^13^C-labeling of *Bacteroidetes* at day 3 as well as the precipitous drop in abundance of *Bacteroidetes* at day 3 followed by ^13^C-labeling of Micrococcales at day 7. Trophic exchange could be enabled by mother cell lysis (in the case of spore formers such as *Paenibacillus*), viral lysis, and/or the direct indirect effects of predation. *Bacteroidetes* types have been shown to become ^13^C-labeled after the addition of live ^13^C-labeled *Escherichia coli* to soil [50] indicating their ability to assimilate C from microbial biomass. In addition, the dominant OTU labeled in the ^13^C-xylose treatment from the *Micrococcales* shares 100% SSU rRNA gene sequence identity to *Agromyces ramosus* a known predator that feeds upon on many microorganisms including yeast and *Micrococcus luteus* [51]. *Agromyces* are abundant microorganisms in many soils and *Agromyces ramosus* was the most abundant xylose responder in our experiment – the fourth most abundant OTU in our dataset. It is notable however, that if *Agromyces ramosus* is acting as a predator in our experiment, the organism remains unlabeled in response to ^13^C-cellulose which suggests that its activity may be specific for certain prey or for certain environmental conditions (see SI note 3 for further discussion of trophic C exchange). Climate change is expected to diminish bottom-up controls on microbial growth increasing the importance on top-down biological interactions for mitigating positive climate change feedbacks [47]. Currently the extent of bacterial predatory activity in soil, and its consequences for the soil C-cycle and carbon use efficiency is largely unknown. Elucidating the identities of bacterial predators in soil will assist in assessing the implications of climate change on global soil-C storage.

## Conclusion

Microorganisms govern C-transformations in soil and thereby influence global climate but still we do not know the specific identities of microorganisms that carry out critical C transformations. In this experiment microorganisms from physiologically uncharacterized but cosmopolitan soil lineages participated in cellulose decomposition. Cellulose responders included members of the *Verrucomicrobia (Spartobacteria*), *Chloroflexi, Bacteroidetes* and *Planctomycetes. Spartobacteria* in particular are globally cosmopolitan soil microorganisms and are often the most abundant *Verrucomicrobia* order in soil [49]. Fast-growing aerobic spore formers from *Firmicutes* assimilated labile C in the form of xylose. Xylose responders within the *Bacteroidetes* and *Actinobacteria* likely became labeled by consuming ^13^C-labeled constituents of microbial biomass either by saprotrophy or predation. Our results suggest that cosmopolitan *Spartobacteria* may degrade cellulose on a global scale, decomposition of labile plant C may initiate trophic transfer within the bacterial food web, and life history traits may act as a filter constraining the diversity of active microorganisms relative to those with the genomic potential for a given metabolism.

## Methods

All code to take raw SSU rRNA gene sequencing reads to final publication figures and through all presented analyses is located at the following URL: https://github.com/chuckpr/CSIP_succession_data_analysis.

DNA sequences are deposited on MG-RAST (Accession XXXXXXX).

Twelve soil cores (5 cm diameter × 10 cm depth) were collected from six sampling locations within an organically managed agricultural field in Penn Yan, New York. Soils were sieved (2 mm), homogenized, distributed into flasks (10 g in each 250 ml flask, n = 36) and equilibrated for 2 weeks. We amended soils with a mixture containing 2.9 mg C g^−1^ soil dry weight (d.w.) and brought soil to 50% water holding capacity. By mass the amendment contained 38% cellulose, 23% lignin, 20% xylose, 3% arabinose, 1% galactose, 1% glucose, and 0.5% mannose. 10.6% amino acids (made in house based on Teknova C9795 formulation) and 2.9% Murashige Skoog basal salt mixture which contains macro and micro-nutrients that are associated with plant biomass (Sigma Aldrich M5524). This mixture approximates the molecular composition of switchgrass biomass with hemicellulose replaced by its constituent monomers [52]. We set up three parallel treatments varying the isotopically labeled component in each treatment. The treatments were (1) a control treatment with all unlabeled components, (2) a treatment with ^13^C-cellulose instead of unlabeled cellulose (synthesized as described in SI), and (3) a treatment with ^13^C-xylose (98 atom% ^13^C, Sigma Aldrich) instead of unlabeled xylose. Other details relating to substrate addition can be found in SI. Microcosms were sampled destructively at days 1 (control and xylose only), 3, 7, 14, and 30 and soils were stored at −80° C until nucleic acid extraction. The abbreviation 13CXPS refers to the ^13^C-xylose treatment (^13^C Xylose Plant Simulant), 13CCPS refers to the ^13^C-cellulose treatment, and 12CCPS refers to the control treatment.

We used DESeq2 (R package), an RNA-Seq differential expression statistical framework [53], to identify OTUs that were enriched in high density gradient fractions from ^13^C-treatments relative to corresponding gradient fractions from control treatments (for review of RNA-Seq differential expression statistics applied to microbiome OTU count data see [54]). We define “high density gradient fractions” as gradient fractions whose density falls between 1.7125 and 1.755 g ml^−1^. For each OTU, we calculates logarithmic fold change (LFC) and corresponding standard error for enrichment in high density fractions of ^13^C treatments relative to control. Subsequently, a one-sided Wald test was used to assess the statistical significance of LFC values with the null hypothesis that LFC was less than one standard deviation above the mean of all LFC values. We independently filtered OTUs prior to multiple comparison corrections on the basis of sparsity eliminating OTUs that failed to appear in at least 45% of high density fractions for a given comparison. P-values were adjusted for multiple comparisons using the Benjamini and Hochberg method [55]. We selected a false discovery rate of 10% to denote statistical significance.

See SI for additional information on experimental and analytical methods.

## Acknowledgements

The authors would like to acknowledge the assistance of John Christian Gaby and Mallory Choudoir in developing the method used to produce ^13^C-labeled cellulose. We would also like to thank Ruth Ley, Steve Zinder, Nelson Hairston, and Nick Youngblut for providing comments that were helpful in the development of this manuscript. Sander Hunter assisted with the microcosm set up. This material is based upon work supported by the Department of Energy Office of Science, Office of Biological & Environmental Research Genomic Science Program under Award Numbers DE-SC0004486 and DE-SC0010558. Manuscript preparation by Ashley N. Campbell was performed under the auspices of the U.S. Department of Energy by LLNL under Contract DE-AC52-07NA27344.

## Supplemental Figures and Tables

**Fig. S1.**
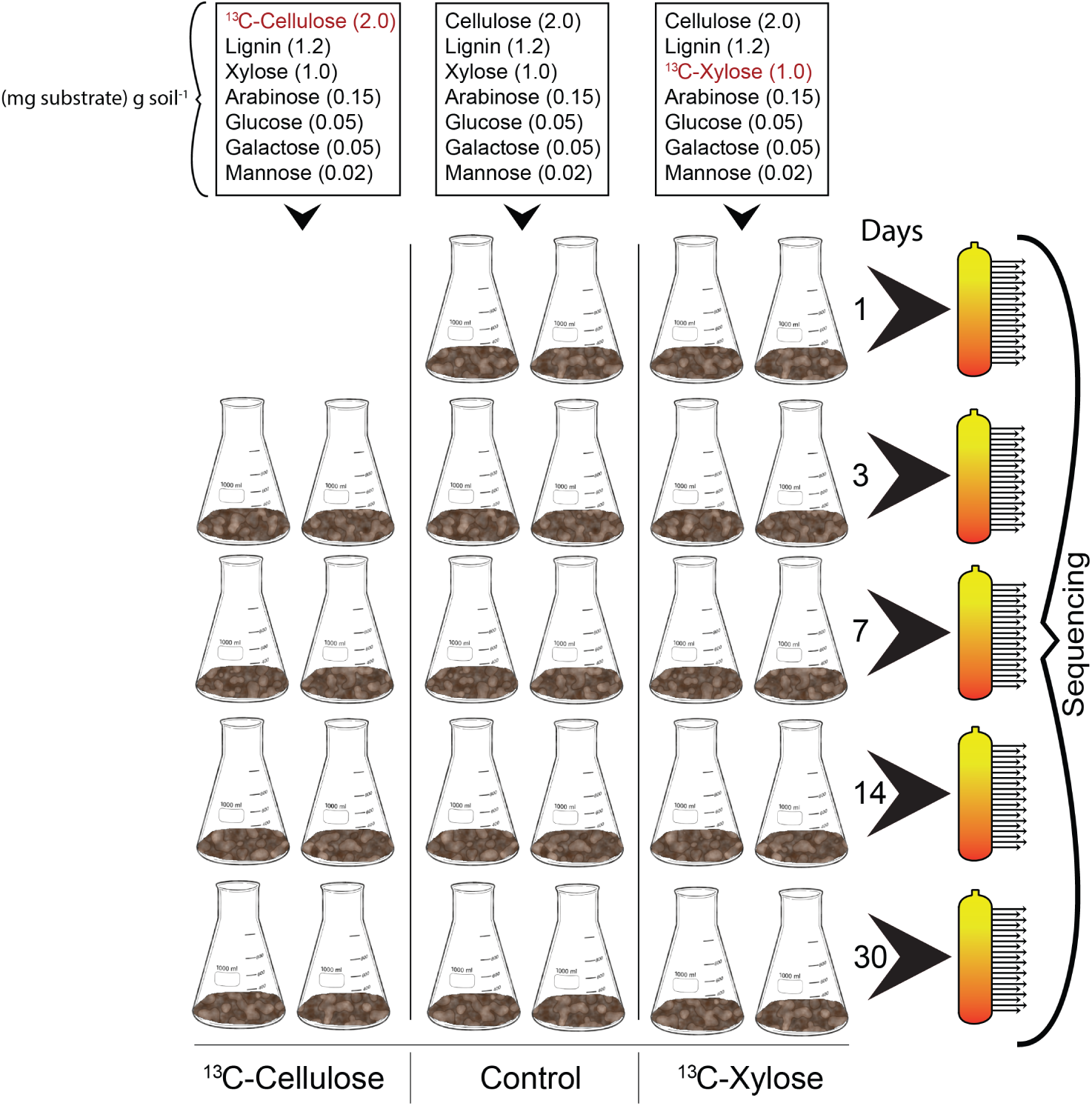
We added a carbon mixture that contained inorganic salts and amino acids (not shown here) to each soil microcosm where the only difference between treatments was the ^13^C-labeled isotope (in red). At days 1, 3, 7, 14, and 30 replicate microcosms were destructively harvested for downstream molecular applications. DNA from each treatment and time (n = 14) was subjected to CsCl density gradient centrifugation and density gradients were fractionated (orange tubes wherein each arrow represents a fraction from the density gradient). SSU rRNA genes from each gradient fraction were PCR amplified and sequenced. In addition, SSU rRNA genes were also PCR amplified and sequenced from non-fractionated DNA to represent the soil microbial community.

**Fig. S2.**
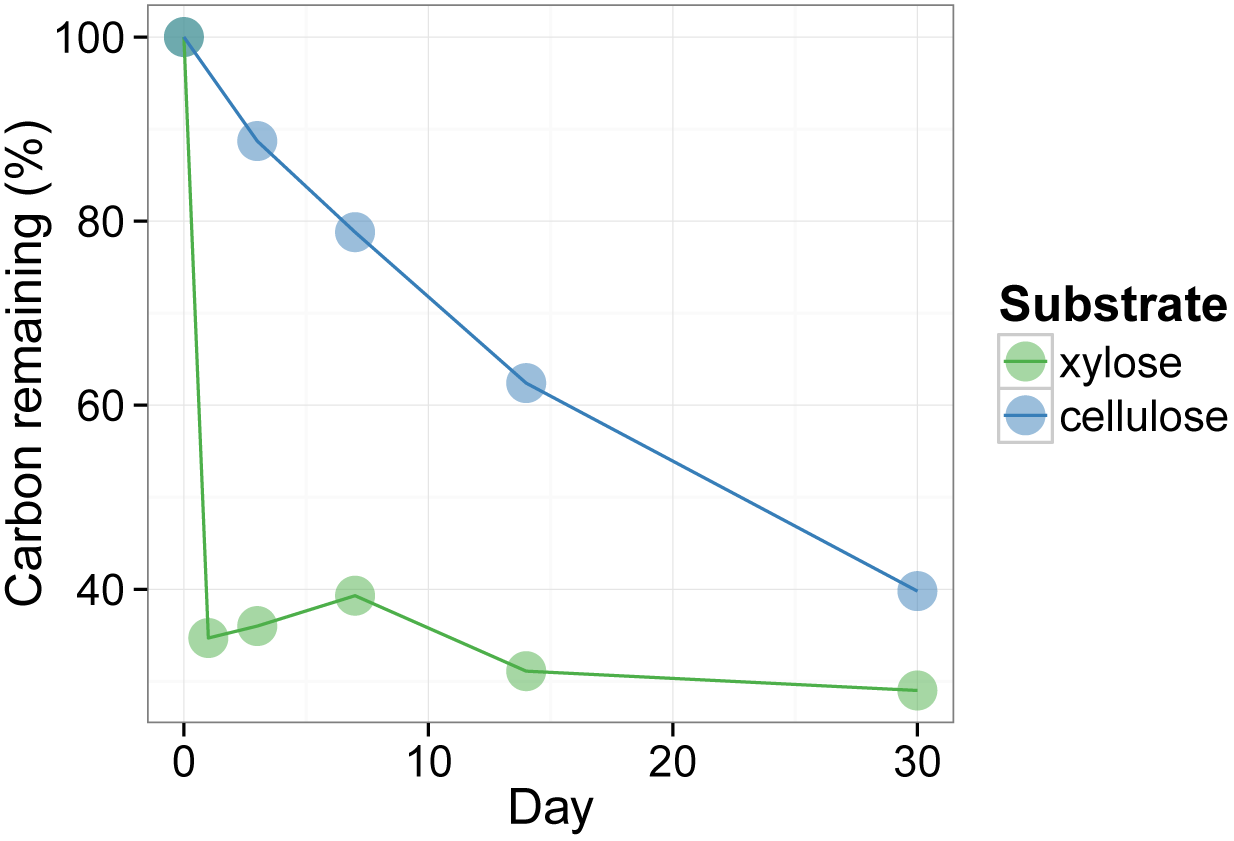
The metabolization of ^13^C-xylose and ^13^C-cellulose is indicated by the percentage of the added ^13^C that remains in soil over time.

**Fig. S3.**
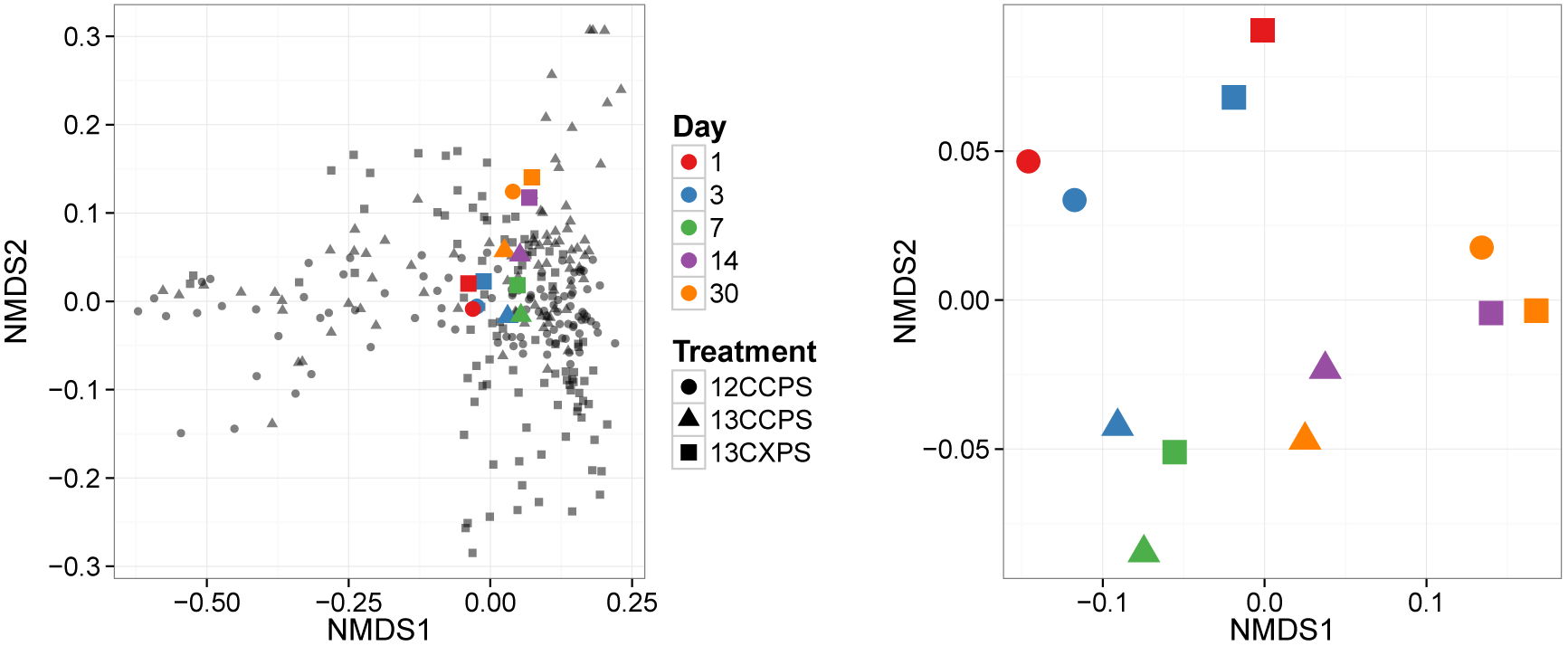
NMDS analysis of SSU rRNA gene composition in non-fractionated DNA (colored points) indicates that isotopic labelling does not alter overall microbial community composition, microbial community composition in the soil microcosms changes over time, and variance in non-fractionated DNA is smaller than variance in fractionated DNA (black points). SSU rRNA gene sequences were determined for non-fractionated DNA from the unlabeled control, ^13^C-xylose, and ^13^C-cellulose treatments over time (colors indicate time, different symbols used for different treatments). Distance in SSU rRNA gene composition was quantified with the UniFrac metric. The leftmost panel indicates NMDS of data from both non-fractionated and fractionated samples. The rightmost panel indicates NMDS of data only from non-fractionated DNA. Statistical analysis is presented in main text.

**Fig. S4.**
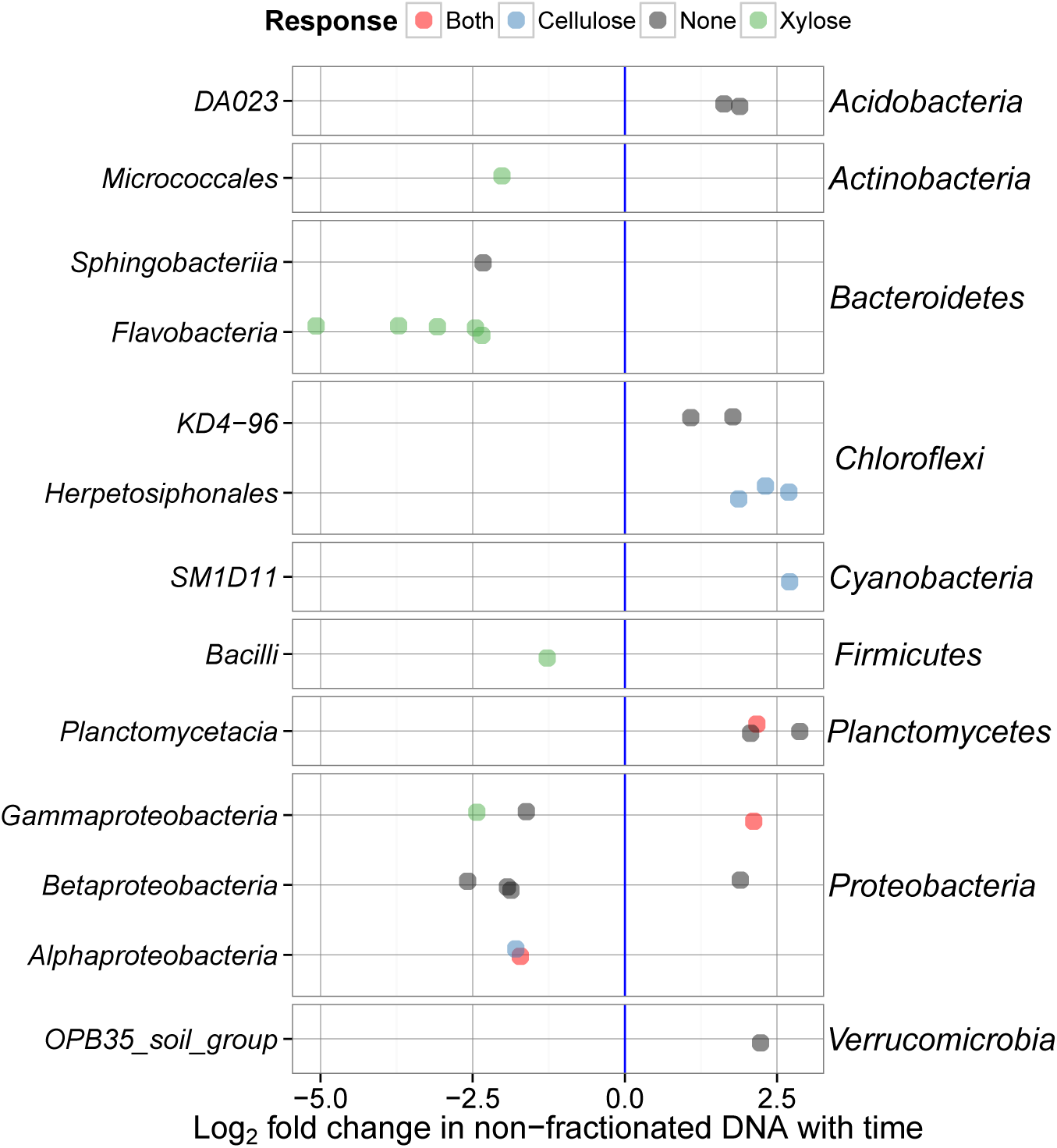
Change in non-fractionated DNA relative abundance versus time (expressed as LFC) for OTUs that changed significantly over time (P-value < 0.10, Wald test). Each panel shows one phylum (labeled on the right). The taxonomic class is indicated on the left. Colors represent results shown in and. OTUs that responded to just xylose are shown in green, just cellulose in blue, and both xylose and cellulose in red.

**Fig. S5.**
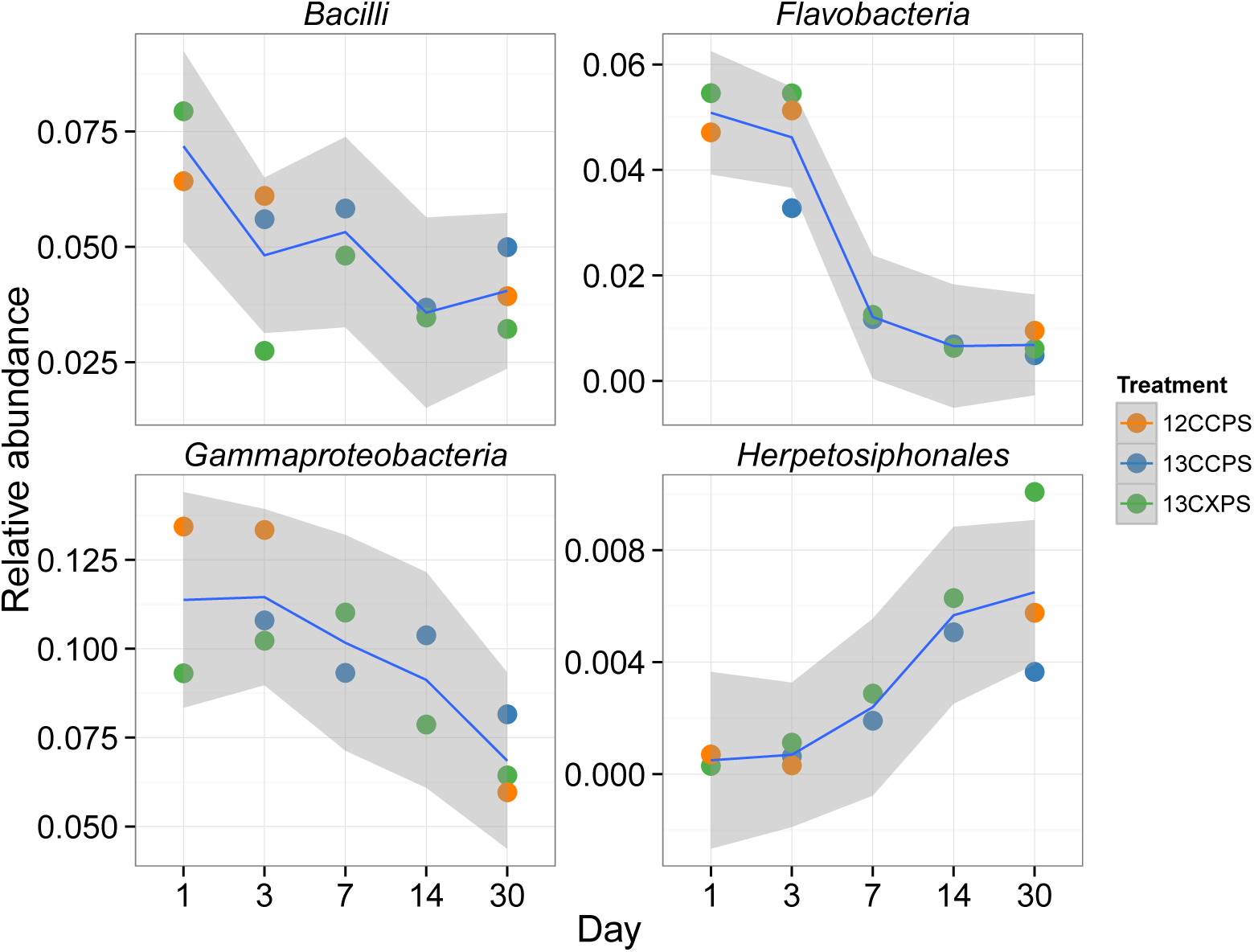
Relative abundance in non-fractionated DNA versus time for classes that changed significantly. Samples from different treatments are labeled with different colors as indicated in the scale. Statistical analysis is presented in main text.

**Fig. S6.**
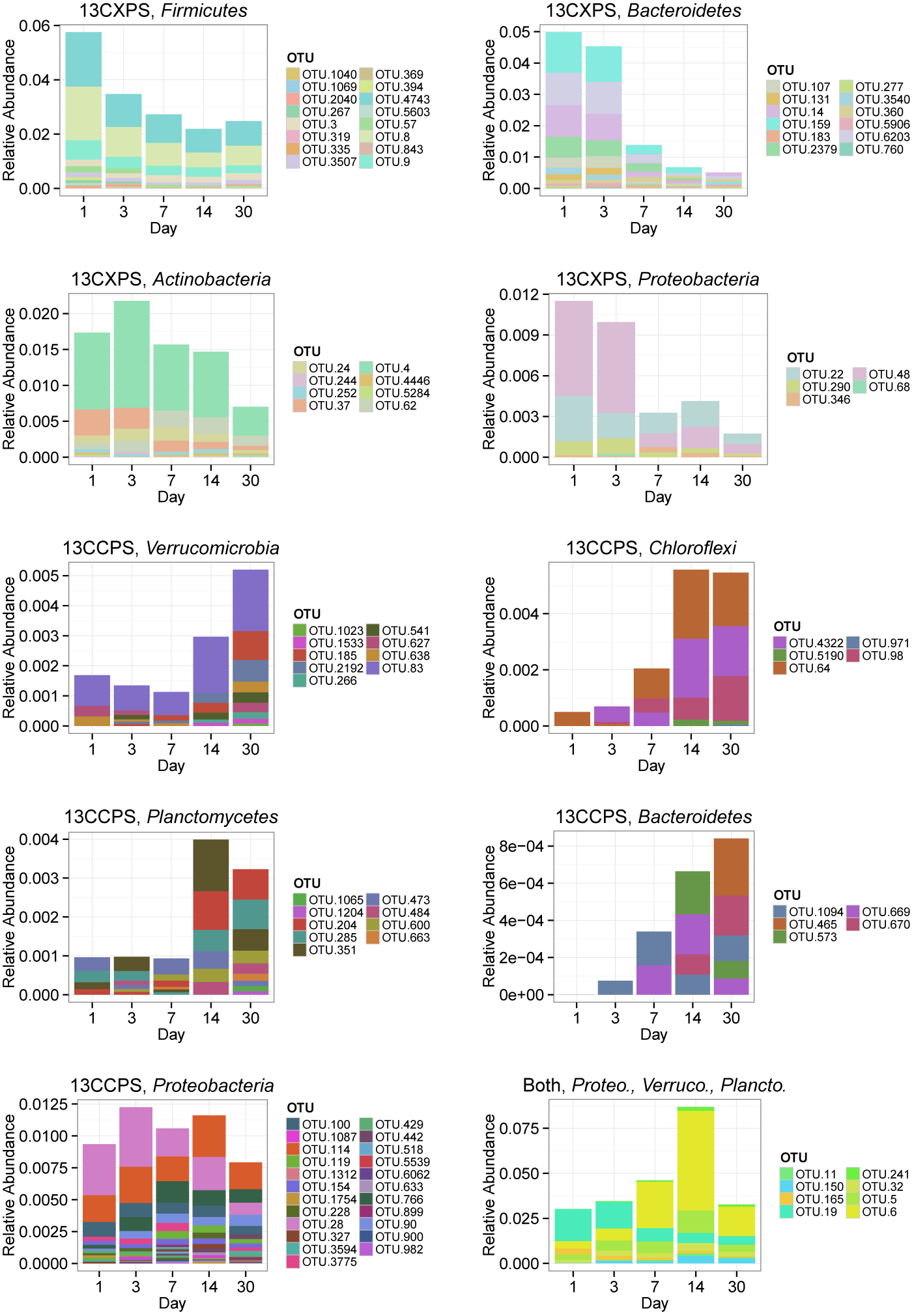
Change in relative abundance in non-fractionated DNA over time for xylose responders (13CXPS) and cellulose responders (13CCPS). Each panel represents a responders to the indicated substrate (i.e. cellulose (13CCPS) or xylose (13CXPS)) within the indicated phylum except for the lower right panel which shows all reponders to both xylose and celluose. The abbreviations Proteo., Verruco., and Plancto., correspond to *Proteobacteria, Verrucomicrobia,* and *Planctomycetes,* respectively.

**Fig. S7.**
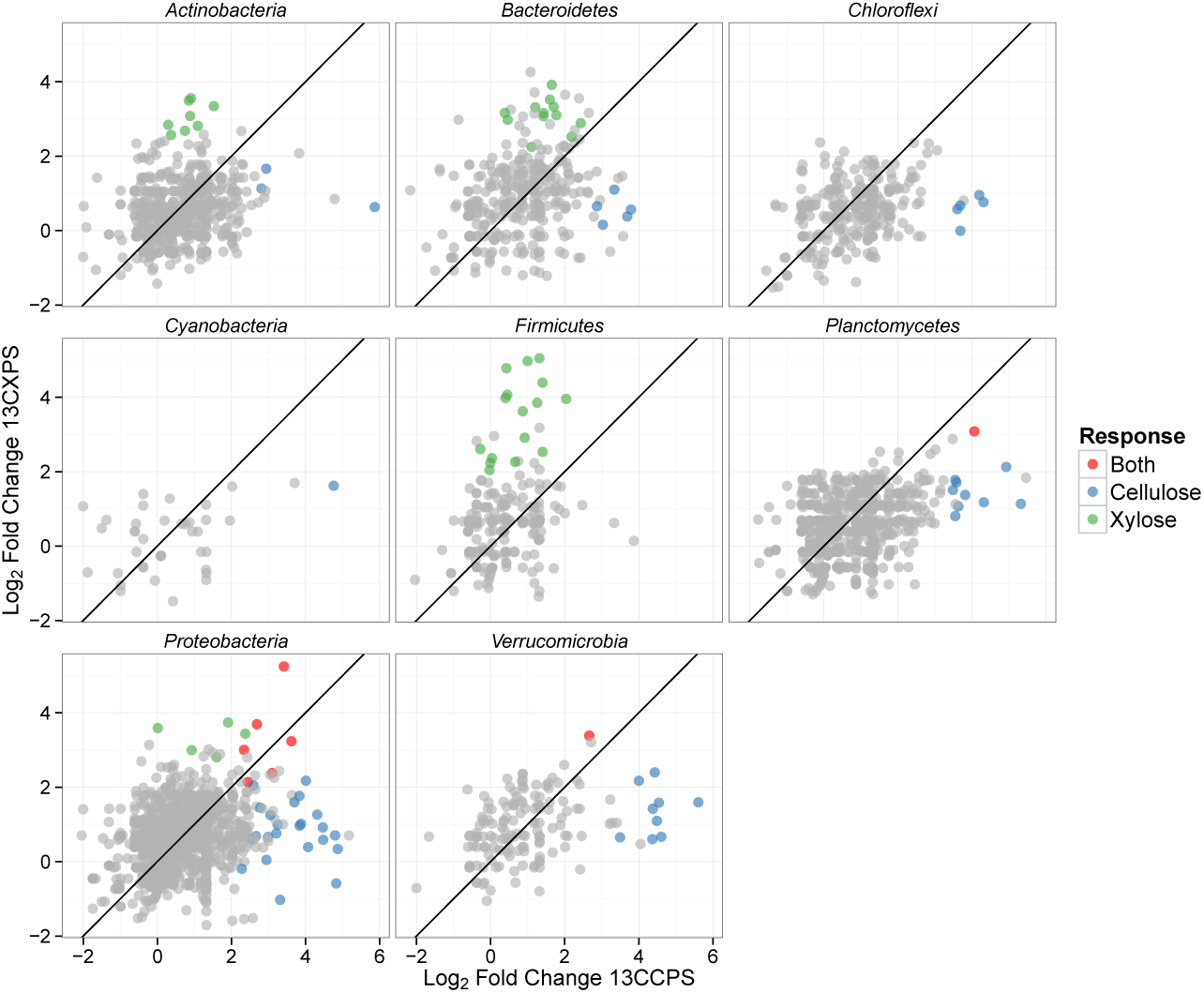
Maximum enrichment at any point in time in high density fractions of ^13^C-treatments relative to control (expressed as LFC) shown for ^13^C-cellulose versus ^13^C-xylose treatments. Each point represents an OTU. Blue points are cellulose responders, green xylose responders, red are responders to both xylose and cellulose, and gray points are OTUs that did not respond to either substrate. Line indicates a slope of one.

**Fig. S8.**
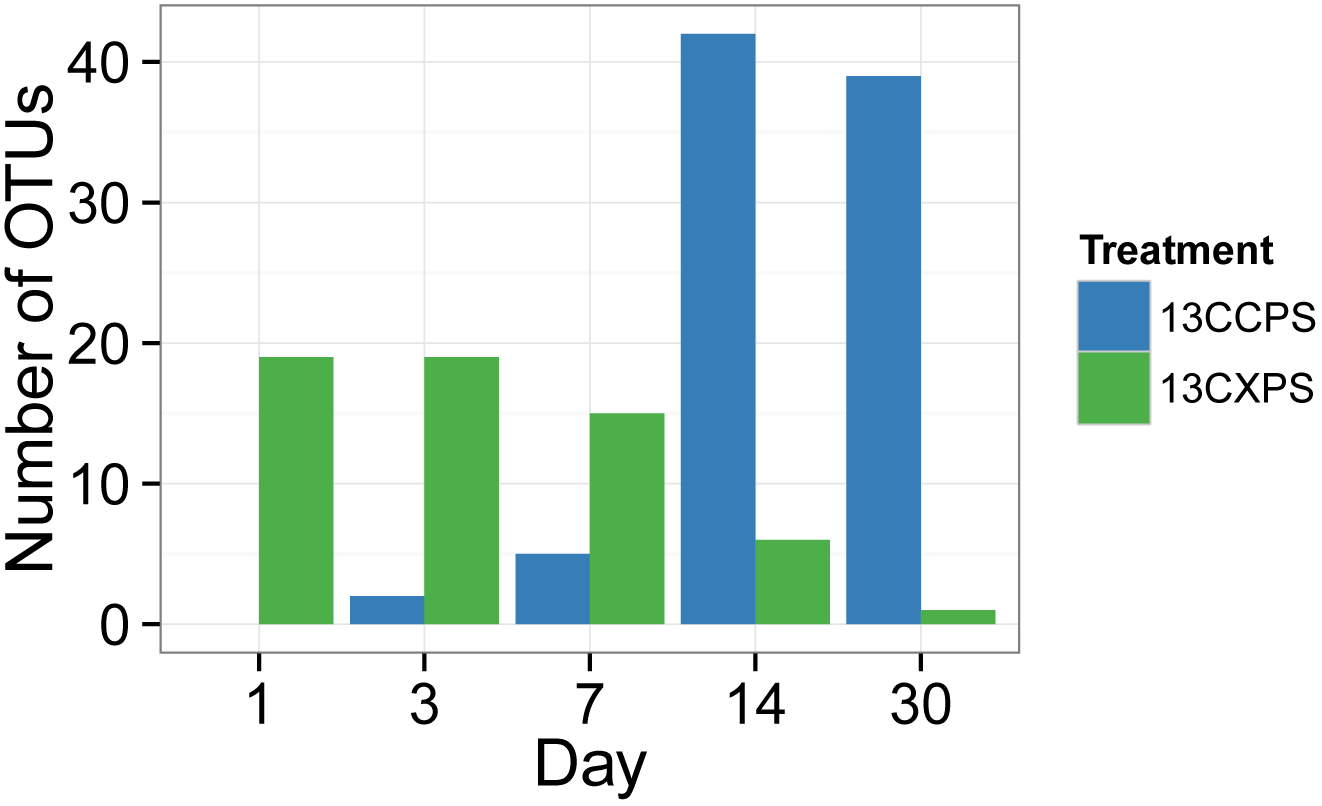
Counts of xylose responders and cellulose responders over time.

**Fig. S9.**
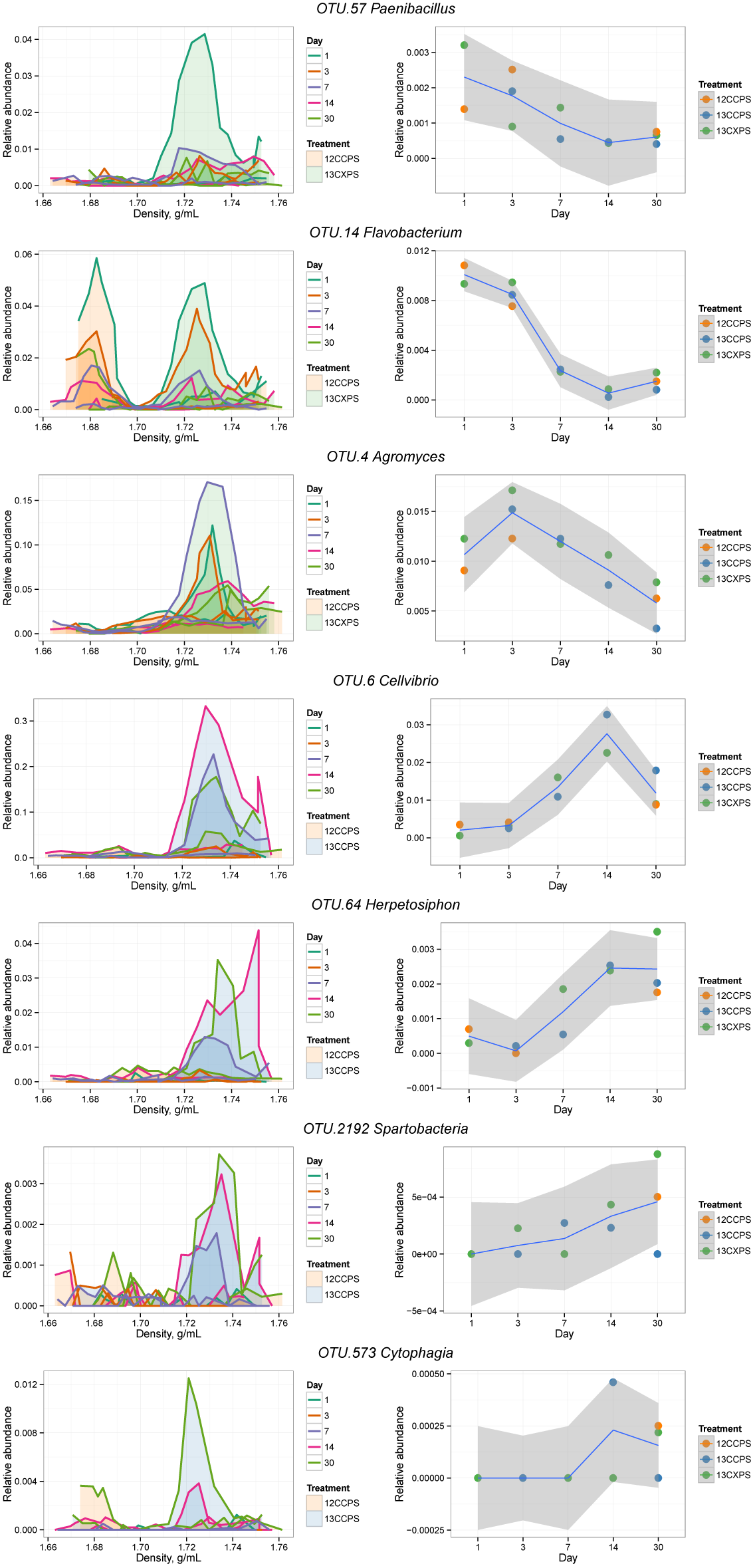
Raw data from individual responders highlighted in the main text (see Results). The left column shows OTU relative abundance in density gradient fractions for the indicated treatment pair at each sampling point. Time is indicated by the line color (see legend). Gradient profiles are shaded to represent the different treatments where orange represents “control”, blue “^13^C-cellulose”, and green “^13^C-xylose.” The right column shows the relative abundance of each OTU in non-fractionated DNA. Enrichment in the high density fractions of ^13^C-treatments indicates an OTU likely has ^13^C-labeled DNA.

**Fig. S10.**
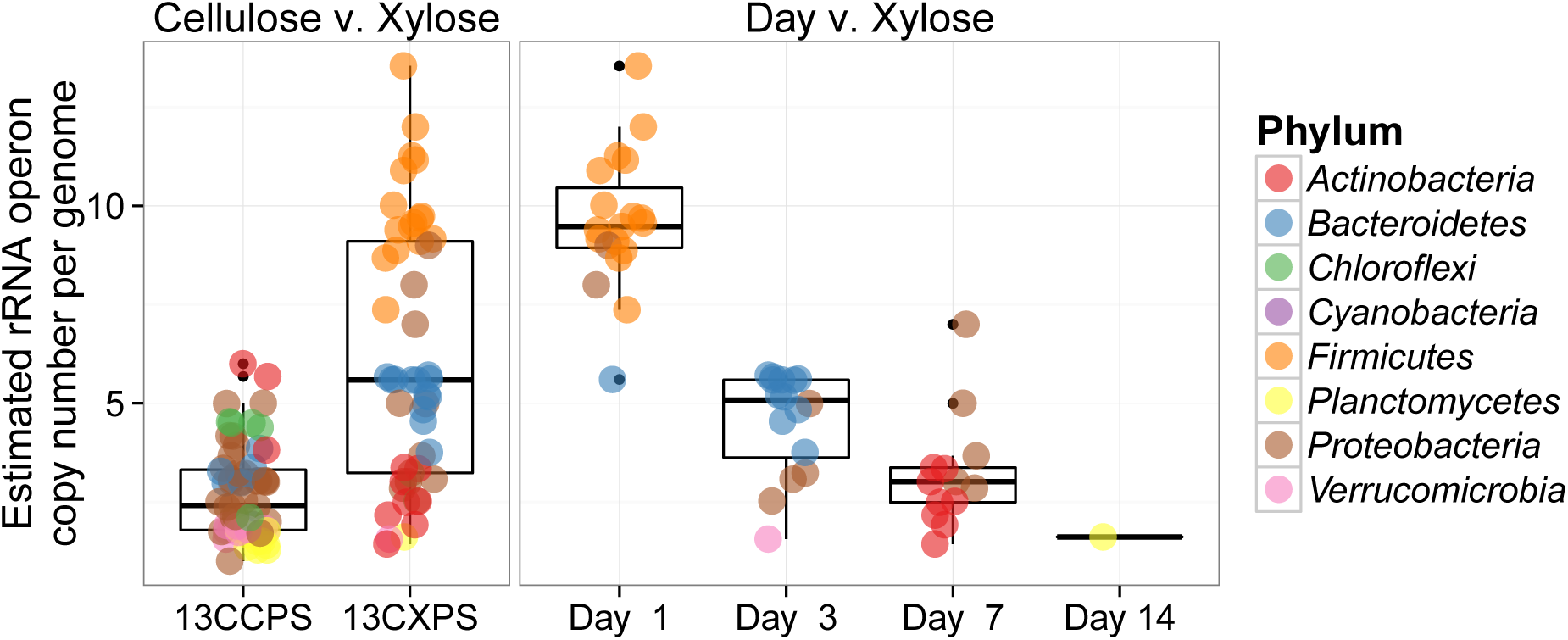
Estimated *rrn* copy number for xylose and cellulose responders. The leftmost panel contrasts estimated *rrn* copy number for cellulose (13CCPS) and xylose (13CXPS) responders. The right panel shows estimated *rrn* copy number versus time of first response for xylose responders. Colors denote the phylum of the OTUs (see legend).

**Fig. S11.**
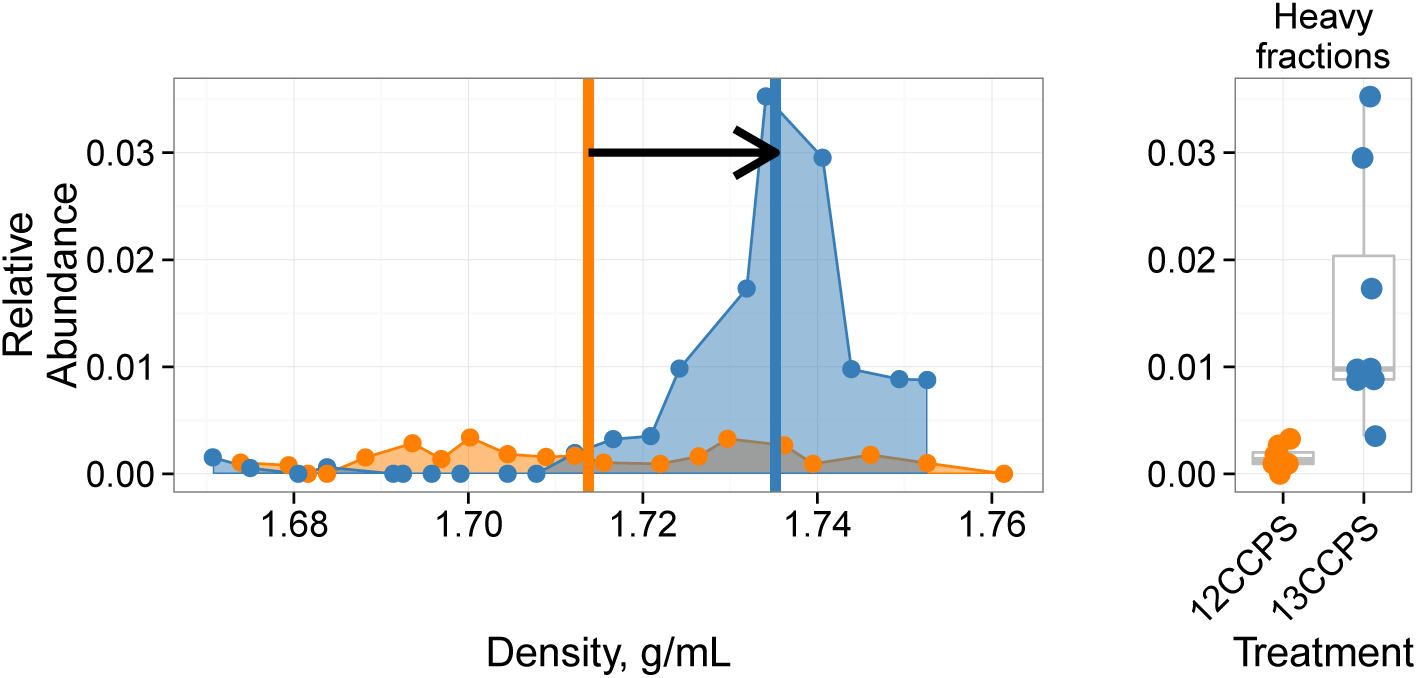
Density profile for a single cellulose responder in the ^13^C-cellulose treatment (blue) and control (orange). Vertical lines show center of mass for each density profile and the arrow denotes the magnitude and direction of 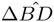. Right panel shows relative abundance values in the high density fractions (The boxplot line is the median value. The box spans one interquartile range (IR) about the median, whiskers extend 1.5 times the IR, and the dots indicate outlier values beyond 1.5 times the IR).

**Fig. S12.**
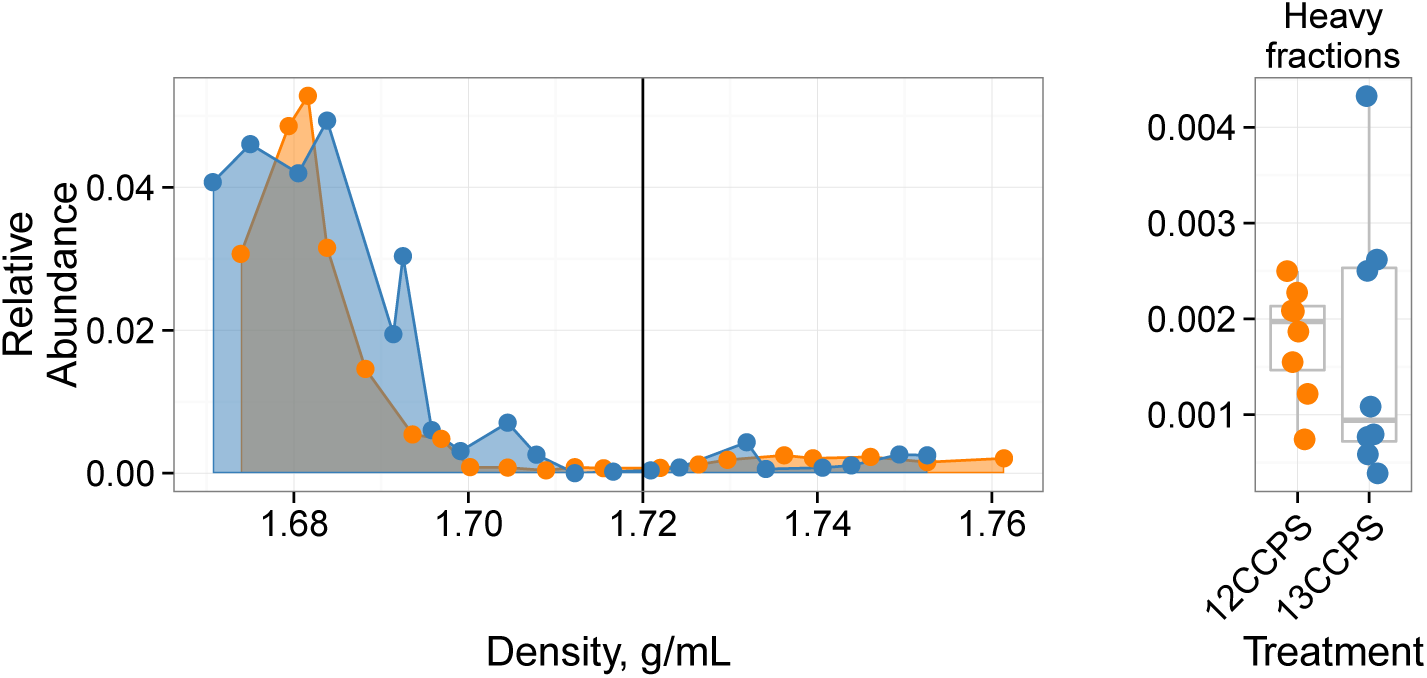
Density profile for a single non-responder OTU. The ^13^C-cellulose treatment is in blue and the control treatment is in orange. The vertical line shows where high density fractions begin as defined in our analysis. The right panel shows relative abundance values in the high density fractions for each gradient (The boxplot line is the median value. The box spans one interquartile range (IR) about the median, whiskers extend 1.5 times the IR and the dots indicate outlier values beyond 1.5 times the IR).

**Table S1:** ^13^C-xylose responders BLAST against Living Tree Project

| OTU ID | Fold change <sup>a</sup> | Day <sup>b</sup> | All days <sup>c</sup> | Top BLAST hits | BLAST %ID | Phylum;Class;Order |
| --- | --- | --- | --- | --- | --- | --- |
| OTU.1040 | 4.78 | 1 | 1 | *Paenibacillus daejeonensis* | 100.0 | Firmicutes Bacilli Bacillales |
| OTU.1069 | 3.85 | 1 | 1 | *Paenibacillus terrigena* | 100.0 | Firmicutes Bacilli Bacillales |
| OTU.107 | 2.25 | 3 | 3 | *Flavobacterium* sp. 15C3,<br>*Flavobacterium banpakuense* | 99.54 | Bacteroidetes Flavobacteria<br>Flavobacteriales |
| OTU.11 | 5.25 | 7 | 7 | *Stenotrophomonas pavanii*,<br>*Stenotrophomonas maltophilia*,<br>*Pseudomonas geniculata* | 99.54 | Proteobacteria<br>Gammaproteobacteria<br>Xanthomonadales |
| OTU.131 | 3.07 | 3 | 3 | *Flavobacterium fluvii*,<br>*Flavobacteria bacterium HMD1033*,<br>*Flavobacterium* sp. HMD1001 | 100.0 | Bacteroidetes Flavobacteria<br>Flavobacteriales |
| OTU.14 | 3.92 | 3 | 1, 3 | *Flavobacterium oncorhynchi*,<br>*Flavobacterium glycines*,<br>*Flavobacterium succinicans* | 99.09 | Bacteroidetes Flavobacteria<br>Flavobacteriales |
| OTU.150 | 3.08 | 14 | 14 | No hits of at least 90%<br>identity | 86.76 | Planctomycetes Planctomycetacia<br>Planctomycetales |
| OTU.159 | 3.16 | 3 | 3 | *Flavobacterium hibernum* | 98.17 | Bacteroidetes Flavobacteria<br>Flavobacteriales |
| OTU.165 | 2.38 | 3 | 3 | *Rhizobium skierniewicense*,<br>*Rhizobium vignae*,<br>*Rhizobium larrymoorei*,<br>*Rhizobium alkalisoli*,<br>*Rhizobium galegae*,<br>*Rhizobium huautlense* | 100.0 | Proteobacteria Alphaproteobacteria<br>Rhizobiales |
| OTU.183 | 3.31 | 3 | 3 | No hits of at least 90%<br>identity | 89.5 | Bacteroidetes Sphingobacteriia<br>Sphingobacteriales |
| OTU.19 | 2.14 | 7 | 7 | *Rhizobium alamii*,<br>*Rhizobium mesosinicum*,<br>*Rhizobium mongolense*,<br>*Arthrobacter viscosus*,<br>*Rhizobium sullae*,<br>*Rhizobium yanglingense*,<br>*Rhizobium loessense* | 99.54 | Proteobacteria Alphaproteobacteria<br>Rhizobiales |
| OTU.2040 | 2.91 | 1 | 1 | *Paenibacillus pectinilyticus* | 100.0 | Firmicutes Bacilli Bacillales |
| OTU.22 | 2.8 | 7 | 7, 14 | *Paracoccus* sp. NB88 | 99.09 | Proteobacteria Alphaproteobacteria<br>Rhodobacterales |
| OTU.2379 | 3.1 | 3 | 3 | *Flavobacterium pectinovorum*,<br>*Flavobacterium* sp. CS100 | 97.72 | Bacteroidetes Flavobacteria<br>Flavobacteriales |
| OTU.24 | 2.81 | 7 | 7 | *Cellulomonas aerilata*,<br>*Cellulomonas humilata*,<br>*Cellulomonas terrae*,<br>*Cellulomonas soli*,<br>*Cellulomonas xylanilytica* | 100.0 | Actinobacteria Micrococcales<br>Cellulomonadaceae |
| OTU.241 | 3.38 | 3 | 3, 14 | No hits of at least 90%<br>identity | 87.73 | Verrucomicrobia Spartobacteria<br>Chthoniobacteriales |
| OTU.244 | 3.08 | 7 | 7 | *Cellulosimicrobium funkei*,<br>*Cellulosimicrobium terreum* | 100.0 | Actinobacteria Micrococcales<br>Promicromonosporaceae |
| OTU.252 | 3.34 | 7 | 7 | *Promicromonospora thailandica* | 100.0 | Actinobacteria Micrococcales<br>Promicromonosporaceae |
| OTU.267 | 4.97 | 1 | 1 | *Paenibacillus pabuli*,<br>*Paenibacillus tundrae*,<br>*Paenibacillus taichungensis*,<br>*Paenibacillus xylanexedens*,<br>*Paenibacillus xylanilyticus* | 100.0 | Firmicutes Bacilli Bacillales |
| OTU.277 | 3.52 | 3 | 3 | *Solibius ginsengiterrae* | 95.43 | Bacteroidetes Sphingobacteriia<br>Sphingobacteriales |

Table S1 – continued from previous page
| OTU ID | Fold change | Day | All days | Top BLAST hits | BLAST %ID | Phylum;Class;Order |
| --- | --- | --- | --- | --- | --- | --- |
| OTU.290 | 3.59 | 1 | 1 | *Pantoea* spp.,<br>*Kluyvera* spp.,<br>*Klebsiella* spp.,<br>*Erwinia* spp.,<br>*Enterobacter* spp.,<br>*Buttiauxella* spp. | 100.0 | *Proteobacteria*<br>*Gammaproteobacteria*<br>*Enterobacteriales* |
| OTU.3 | 2.61 | 1 | 1 | *[Brevibacterium] frigoritolerans*,<br>*Bacillus* sp. LMG 20238,<br>*Bacillus coahuilensis* m4-4,<br>*Bacillus simplex* | 100.0 | *Firmicutes* *Bacilli* *Bacillales* |
| OTU.319 | 3.98 | 1 | 1 | *Paenibacillus xinjiangensis* | 97.25 | *Firmicutes* *Bacilli* *Bacillales* |
| OTU.32 | 3.0 | 3 | 3, 7, 14 | *Sandaracinus amylolyticus* | 94.98 | *Proteobacteria* *Deltaproteobacteria*<br>*Myxococcales* |
| OTU.335 | 2.53 | 1 | 1 | *Paenibacillus thailandensis* | 98.17 | *Firmicutes* *Bacilli* *Bacillales* |
| OTU.346 | 3.44 | 3 | 3 | *Pseudoduganella violaceinigra* | 99.54 | *Proteobacteria* *Betaproteobacteria*<br>*Burkholderiales* |
| OTU.3507 | 2.36 | 1 | 1 | *Bacillus* spp. | 98.63 | *Firmicutes* *Bacilli* *Bacillales* |
| OTU.3540 | 2.52 | 3 | 3 | *Flavobacterium terrigena* | 99.54 | *Bacteroidetes* *Flavobacteria*<br>*Flavobacteriales* |
| OTU.360 | 2.98 | 3 | 3 | *Flavisolibacter ginsengisoli* | 95.0 | *Bacteroidetes* *Sphingobacteriia*<br>*Sphingobacteriales* |
| OTU.369 | 5.05 | 1 | 1 | *Paenibacillus* sp. D75,<br>*Paenibacillus glycanilyticus* | 100.0 | *Firmicutes* *Bacilli* *Bacillales* |
| OTU.37 | 2.68 | 7 | 7 | *Phycicola gilvus*,<br>*Microterricola viridarii*,<br>*Frigoribacterium faeni*,<br>*Fronidihabitans* sp. RS-15,<br>*Fronidihabitans australicus* | 100.0 | *Actinobacteria* *Micrococcales*<br>*Microbacteriaceae* |
| OTU.394 | 4.06 | 1 | 1 | *Paenibacillus pocheonensis* | 100.0 | *Firmicutes* *Bacilli* *Bacillales* |
| OTU.4 | 2.84 | 7 | 7, 14 | *Agromyces ramosus* | 100.0 | *Actinobacteria* *Micrococcales*<br>*Microbacteriaceae* |
| OTU.4446 | 3.49 | 7 | 7 | *Catenuloplanes niger*,<br>*Catenuloplanes castaneus*,<br>*Catenuloplanes atrovinosus*,<br>*Catenuloplanes crispus*,<br>*Catenuloplanes nepalensis*,<br>*Catenuloplanes japonicus* | 97.72 | *Actinobacteria* *Frankiales*<br>*Nakamurellaceae* |
| OTU.4743 | 2.24 | 1 | 1 | *Lysinibacillus fusiformis*,<br>*Lysinibacillus sphaericus* | 99.09 | *Firmicutes* *Bacilli* *Bacillales* |
| OTU.48 | 2.99 | 1 | 1, 3 | *Aeromonas* spp. | 100.0 | *Proteobacteria*<br>*Gammaproteobacteria* *aaa34a10* |
| OTU.5 | 3.69 | 7 | 7 | *Delftia tsuruhatensis*,<br>*Delftia lacustris* | 100.0 | *Proteobacteria* *Betaproteobacteria*<br>*Burkholderiales* |
| OTU.5284 | 3.56 | 7 | 7 | *Isoptericola nanjingensis*,<br>*Isoptericola hypogeus*,<br>*Isoptericola variabilis* | 98.63 | *Actinobacteria* *Micrococcales*<br>*Promicromonosporaceae* |
| OTU.5603 | 3.96 | 1 | 1 | *Paenibacillus uliginis* | 100.0 | *Firmicutes* *Bacilli* *Bacillales* |
| OTU.57 | 4.39 | 1 | 1, 3, 7, 14,<br>30 | *Paenibacillus castaneae* | 98.62 | *Firmicutes* *Bacilli* *Bacillales* |
| OTU.5906 | 3.16 | 3 | 3 | *Terrimonas* sp. M-8 | 96.8 | *Bacteroidetes* *Sphingobacteriia*<br>*Sphingobacteriales* |
| OTU.6 | 3.24 | 3 | 3 | *Cellvibrio fulvus* | 100.0 | *Proteobacteria*<br>*Gammaproteobacteria*<br>*Pseudomonadales* |

Table S1 – continued from previous page
| OTU ID | Fold change | Day | All days | Top BLAST hits | BLAST %ID | Phylum;Class;Order |
| --- | --- | --- | --- | --- | --- | --- |
| OTU.62 | 2.57 | 7 | 7 | *Nakamurella flavida* | 100.0 | *Actinobacteria Frankiales Nakamurellaceae* |
| OTU.6203 | 3.32 | 3 | 3 | *Flavobacterium granuli*,<br>*Flavobacterium glaciei* | 100.0 | *Bacteroidetes Flavobacteria Flavobacteriales* |
| OTU.68 | 3.74 | 7 | 7 | *Shigella flexneri*,<br>*Escherichia fergusonii*,<br>*Escherichia coli*,<br>*Shigella sonnei* | 100.0 | *Proteobacteria Gammaproteobacteria Enterobacteriales* |
| OTU.760 | 2.89 | 3 | 3 | *Dyadobacter hamtensis* | 98.63 | *Bacteroidetes Cytophagia Cytophagales* |
| OTU.8 | 2.26 | 1 | 1 | *Bacillus niacini* | 100.0 | *Firmicutes Bacilli Bacillales* |
| OTU.843 | 3.62 | 1 | 1 | *Paenibacillus agarexedens* | 100.0 | *Firmicutes Bacilli Bacillales* |
| OTU.9 | 2.04 | 1 | 1 | *Bacillus megaterium*,<br>*Bacillus flexus* | 100.0 | *Firmicutes Bacilli Bacillales* |

<sup>a</sup> Maximum observed $\log_2$ of fold change. <sup>b</sup> Day of maximum fold change. <sup>c</sup> All response days.

**Table S2:**
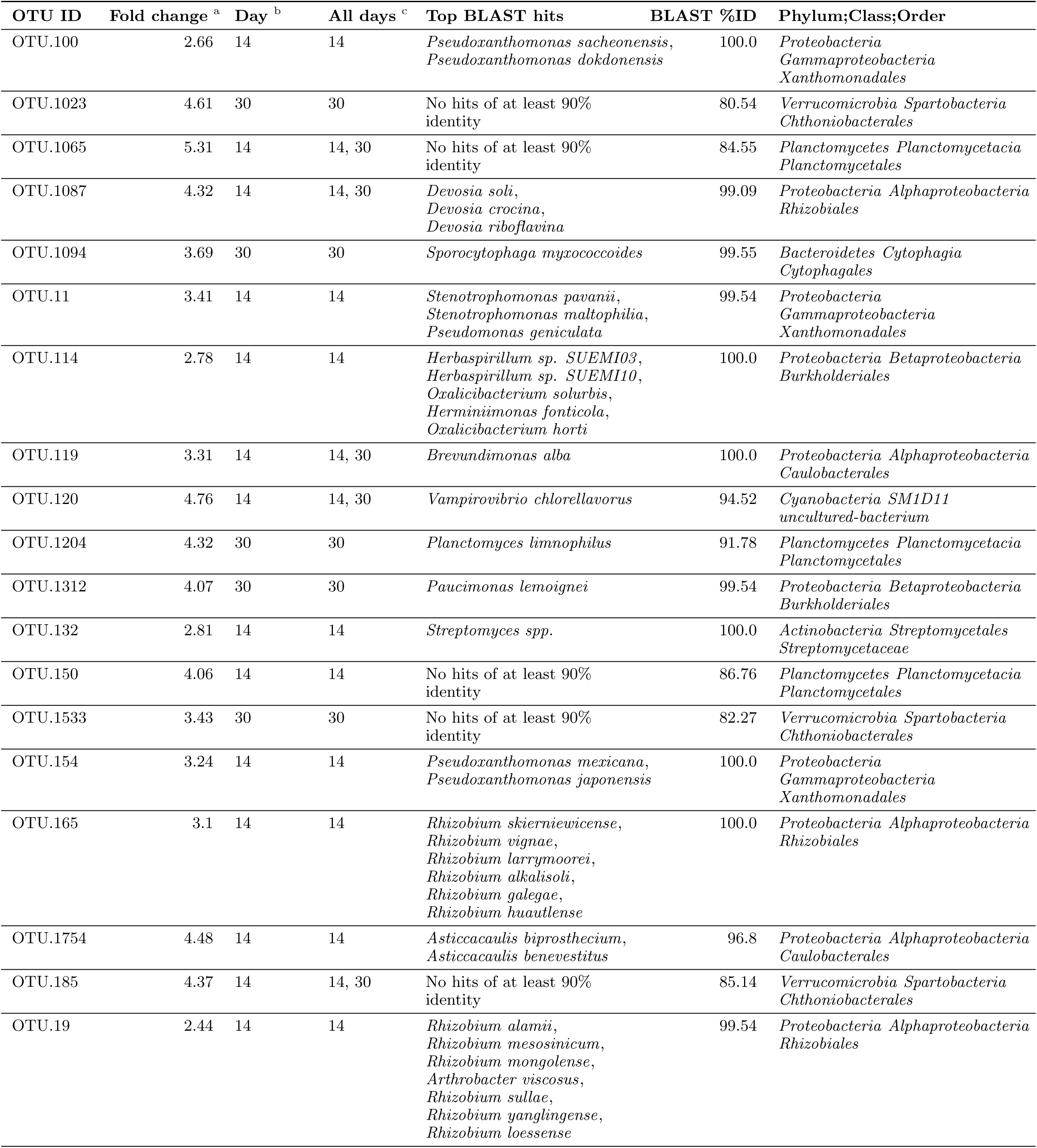
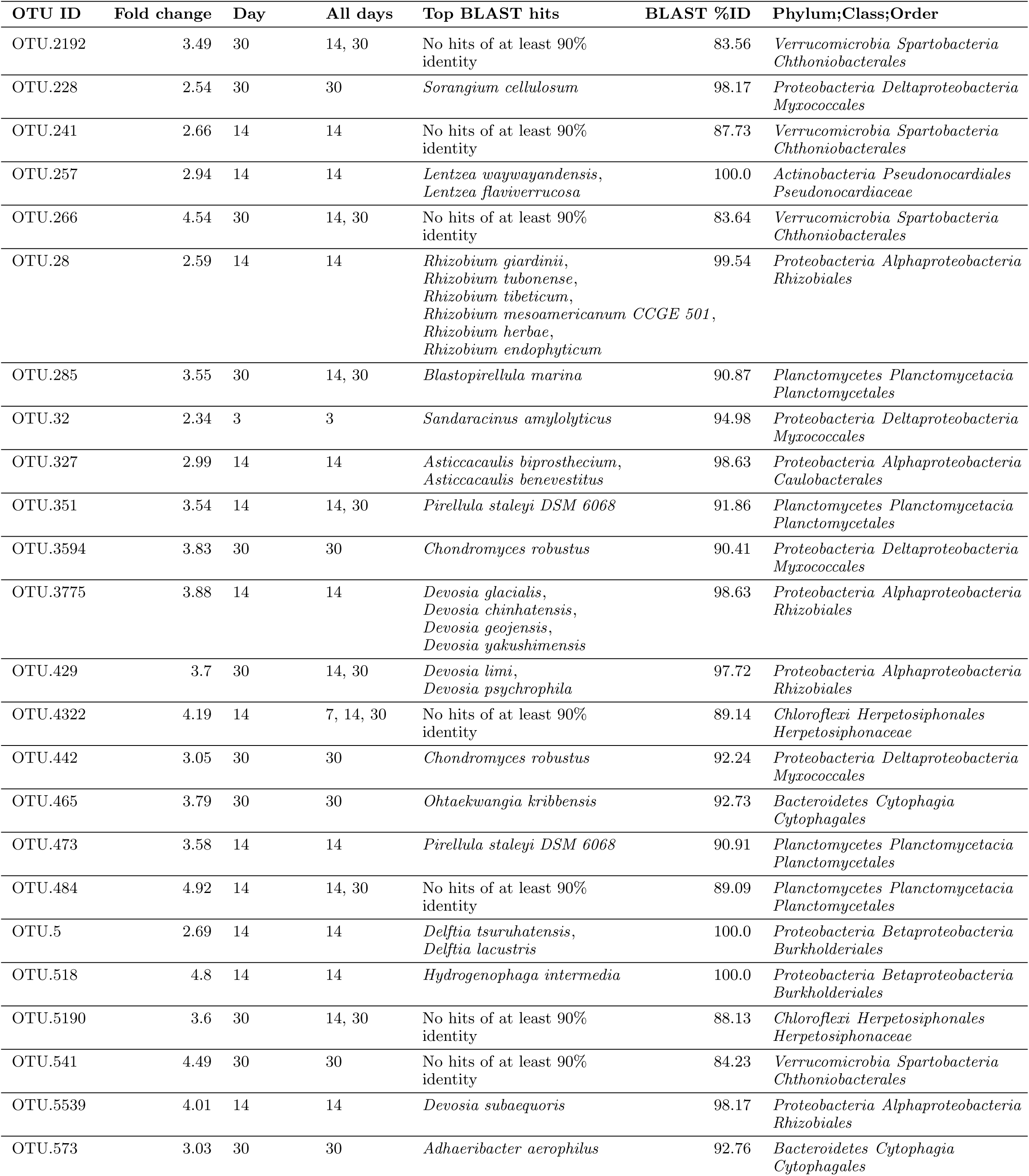
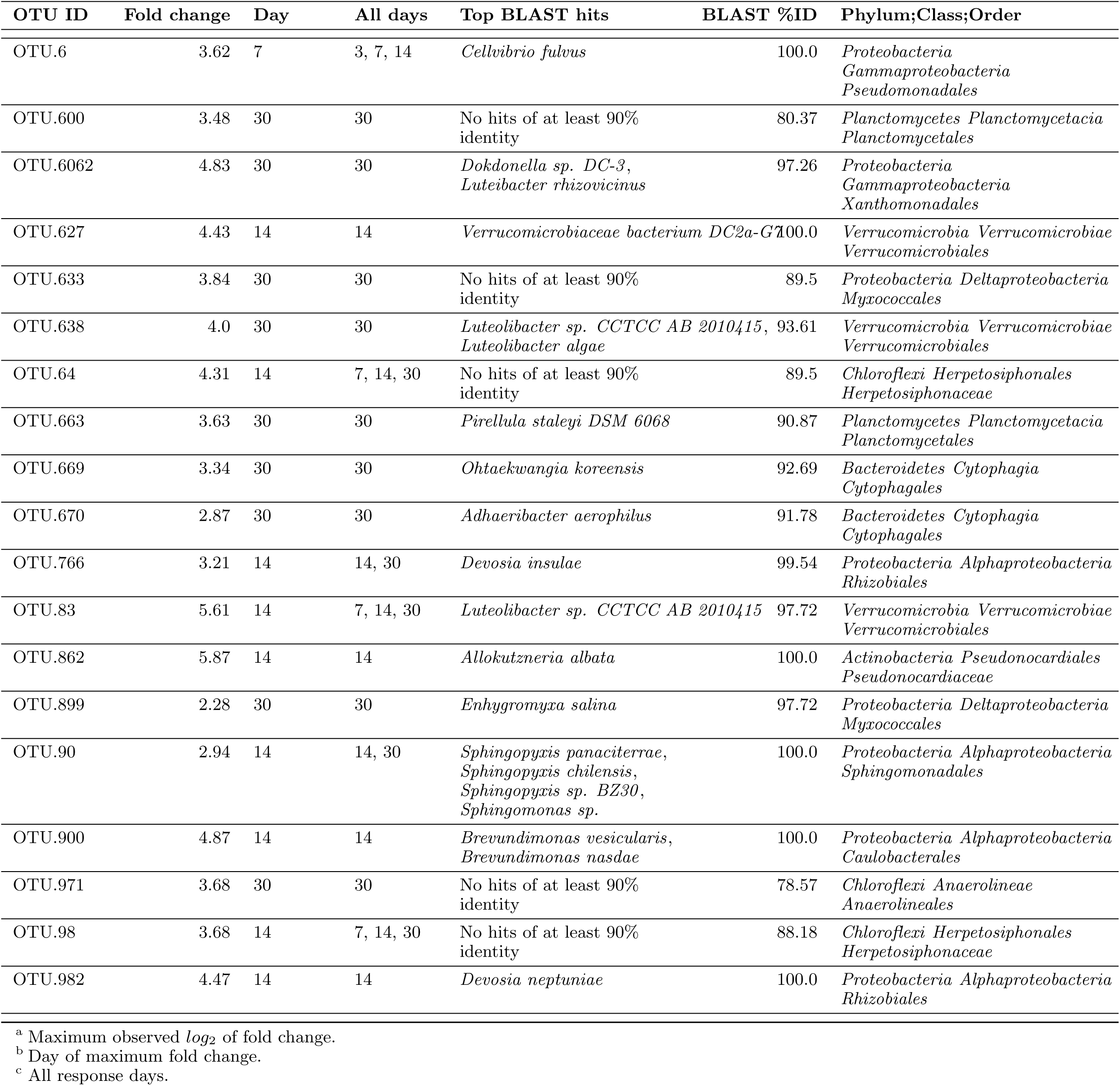
^13^C-cellulose responders BLAST against Living Tree Project

| OTU ID | Fold change <sup>a</sup> | Day <sup>b</sup> | All days <sup>c</sup> | Top BLAST hits | BLAST %ID | Phylum;Class;Order |
| --- | --- | --- | --- | --- | --- | --- |
| OTU.100 | 2.66 | 14 | 14 | *Pseudoxanthomonas sacheonensis*,<br>*Pseudoxanthomonas dokdonensis* | 100.0 | *Proteobacteria*<br>*Gammaproteobacteria*<br>*Xanthomonadales* |
| OTU.1023 | 4.61 | 30 | 30 | No hits of at least 90% identity | 80.54 | *Verrucomicrobia* *Spartobacteria*<br>*Chthoniobacterales* |
| OTU.1065 | 5.31 | 14 | 14, 30 | No hits of at least 90% identity | 84.55 | *Planctomycetes* *Planctomycetacia*<br>*Planctomycetales* |
| OTU.1087 | 4.32 | 14 | 14, 30 | *Devosia soli*,<br>*Devosia crocina*,<br>*Devosia riboflavina* | 99.09 | *Proteobacteria* *Alphaproteobacteria*<br>*Rhizobiales* |
| OTU.1094 | 3.69 | 30 | 30 | *Sporocytophaga myxococcoides* | 99.55 | *Bacteroidetes* *Cytophagia*<br>*Cytophagales* |
| OTU.11 | 3.41 | 14 | 14 | *Stenotrophomonas pavanii*,<br>*Stenotrophomonas maltophilia*,<br>*Pseudomonas geniculata* | 99.54 | *Proteobacteria*<br>*Gammaproteobacteria*<br>*Xanthomonadales* |
| OTU.114 | 2.78 | 14 | 14 | *Herbaspirillum* sp. *SUEMI03*,<br>*Herbaspirillum* sp. *SUEMI10*,<br>*Oxalicibacterium solurbis*,<br>*Hermiinimonas fonticola*,<br>*Oxalicibacterium horti* | 100.0 | *Proteobacteria* *Betaproteobacteria*<br>*Burkholderiales* |
| OTU.119 | 3.31 | 14 | 14, 30 | *Brevundimonas alba* | 100.0 | *Proteobacteria* *Alphaproteobacteria*<br>*Caulobacterales* |
| OTU.120 | 4.76 | 14 | 14, 30 | *Vampirovibrio chlorellavorus* | 94.52 | *Cyanobacteria* *SM1D11*<br>uncultured-bacterium |
| OTU.1204 | 4.32 | 30 | 30 | *Planctomyces limnophilus* | 91.78 | *Planctomycetes* *Planctomycetacia*<br>*Planctomycetales* |
| OTU.1312 | 4.07 | 30 | 30 | *Paucimonas lemoignei* | 99.54 | *Proteobacteria* *Betaproteobacteria*<br>*Burkholderiales* |
| OTU.132 | 2.81 | 14 | 14 | *Streptomyces* spp. | 100.0 | *Actinobacteria* *Streptomycetales*<br>*Streptomycetaceae* |
| OTU.150 | 4.06 | 14 | 14 | No hits of at least 90% identity | 86.76 | *Planctomycetes* *Planctomycetacia*<br>*Planctomycetales* |
| OTU.1533 | 3.43 | 30 | 30 | No hits of at least 90% identity | 82.27 | *Verrucomicrobia* *Spartobacteria*<br>*Chthoniobacterales* |
| OTU.154 | 3.24 | 14 | 14 | *Pseudoxanthomonas mexicana*,<br>*Pseudoxanthomonas japonensis* | 100.0 | *Proteobacteria*<br>*Gammaproteobacteria*<br>*Xanthomonadales* |
| OTU.165 | 3.1 | 14 | 14 | *Rhizobium skierniewicense*,<br>*Rhizobium vignae*,<br>*Rhizobium larrymoorei*,<br>*Rhizobium alkalisoli*,<br>*Rhizobium galegae*,<br>*Rhizobium huautlense* | 100.0 | *Proteobacteria* *Alphaproteobacteria*<br>*Rhizobiales* |
| OTU.1754 | 4.48 | 14 | 14 | *Asticcacaulis biprosthecium*,<br>*Asticcacaulis benevestitus* | 96.8 | *Proteobacteria* *Alphaproteobacteria*<br>*Caulobacterales* |
| OTU.185 | 4.37 | 14 | 14, 30 | No hits of at least 90% identity | 85.14 | *Verrucomicrobia* *Spartobacteria*<br>*Chthoniobacterales* |
| OTU.19 | 2.44 | 14 | 14 | *Rhizobium alamii*,<br>*Rhizobium mesosinicum*,<br>*Rhizobium mongolense*,<br>*Arthrobacter viscosus*,<br>*Rhizobium sullae*,<br>*Rhizobium yanglingense*,<br>*Rhizobium loessense* | 99.54 | *Proteobacteria* *Alphaproteobacteria*<br>*Rhizobiales* |

Table S2 – continued from previous page
| OTU ID | Fold change | Day | All days | Top BLAST hits | BLAST %ID | Phylum;Class;Order |
| --- | --- | --- | --- | --- | --- | --- |
| OTU.2192 | 3.49 | 30 | 14, 30 | No hits of at least 90% identity | 83.56 | Verrucomicrobia Spartobacteria<br>Chthoniobacterales |
| OTU.228 | 2.54 | 30 | 30 | *Sorangium cellulosum* | 98.17 | Proteobacteria Deltaproteobacteria<br>Myxococcales |
| OTU.241 | 2.66 | 14 | 14 | No hits of at least 90% identity | 87.73 | Verrucomicrobia Spartobacteria<br>Chthoniobacterales |
| OTU.257 | 2.94 | 14 | 14 | *Lentzea waywayandensis*,<br>*Lentzea flaviverrucosa* | 100.0 | Actinobacteria Pseudonocardiales<br>Pseudonocardiaceae |
| OTU.266 | 4.54 | 30 | 14, 30 | No hits of at least 90% identity | 83.64 | Verrucomicrobia Spartobacteria<br>Chthoniobacterales |
| OTU.28 | 2.59 | 14 | 14 | *Rhizobium giardinii*,<br>*Rhizobium tubonense*,<br>*Rhizobium tibeticum*,<br>*Rhizobium mesoamericanum* CCGE 501,<br>*Rhizobium herbae*,<br>*Rhizobium endophyticum* | 99.54 | Proteobacteria Alphaproteobacteria<br>Rhizobiales |
| OTU.285 | 3.55 | 30 | 14, 30 | *Blastopirellula marina* | 90.87 | Planctomycetes Planctomycetacia<br>Planctomycetales |
| OTU.32 | 2.34 | 3 | 3 | *Sandaracinus amylolyticus* | 94.98 | Proteobacteria Deltaproteobacteria<br>Myxococcales |
| OTU.327 | 2.99 | 14 | 14 | *Asticcacaulis biprosthecium*,<br>*Asticcacaulis benevestitus* | 98.63 | Proteobacteria Alphaproteobacteria<br>Caulobacterales |
| OTU.351 | 3.54 | 14 | 14, 30 | *Pirellula staleyii* DSM 6068 | 91.86 | Planctomycetes Planctomycetacia<br>Planctomycetales |
| OTU.3594 | 3.83 | 30 | 30 | *Chondromyces robustus* | 90.41 | Proteobacteria Deltaproteobacteria<br>Myxococcales |
| OTU.3775 | 3.88 | 14 | 14 | *Devosia glacialis*,<br>*Devosia chinhatensis*,<br>*Devosia geojensis*,<br>*Devosia yakushimensis* | 98.63 | Proteobacteria Alphaproteobacteria<br>Rhizobiales |
| OTU.429 | 3.7 | 30 | 14, 30 | *Devosia limi*,<br>*Devosia psychrophila* | 97.72 | Proteobacteria Alphaproteobacteria<br>Rhizobiales |
| OTU.4322 | 4.19 | 14 | 7, 14, 30 | No hits of at least 90% identity | 89.14 | Chloroflexi Herpetosiphonales<br>Herpetosiphonaceae |
| OTU.442 | 3.05 | 30 | 30 | *Chondromyces robustus* | 92.24 | Proteobacteria Deltaproteobacteria<br>Myxococcales |
| OTU.465 | 3.79 | 30 | 30 | *Ohtaekwangia kribbensis* | 92.73 | Bacteroidetes Cytophagia<br>Cytophagales |
| OTU.473 | 3.58 | 14 | 14 | *Pirellula staleyii* DSM 6068 | 90.91 | Planctomycetes Planctomycetacia<br>Planctomycetales |
| OTU.484 | 4.92 | 14 | 14, 30 | No hits of at least 90% identity | 89.09 | Planctomycetes Planctomycetacia<br>Planctomycetales |
| OTU.5 | 2.69 | 14 | 14 | *Delftia tsuruhatensis*,<br>*Delftia lacustris* | 100.0 | Proteobacteria Betaproteobacteria<br>Burkholderiales |
| OTU.518 | 4.8 | 14 | 14 | *Hydrogenophaga intermedia* | 100.0 | Proteobacteria Betaproteobacteria<br>Burkholderiales |
| OTU.5190 | 3.6 | 30 | 14, 30 | No hits of at least 90% identity | 88.13 | Chloroflexi Herpetosiphonales<br>Herpetosiphonaceae |
| OTU.541 | 4.49 | 30 | 30 | No hits of at least 90% identity | 84.23 | Verrucomicrobia Spartobacteria<br>Chthoniobacterales |
| OTU.5539 | 4.01 | 14 | 14 | *Devosia subaequoris* | 98.17 | Proteobacteria Alphaproteobacteria<br>Rhizobiales |
| OTU.573 | 3.03 | 30 | 30 | *Adhaeribacter aerophilus* | 92.76 | Bacteroidetes Cytophagia<br>Cytophagales |

Table S2 – continued from previous page
| OTU ID | Fold change | Day | All days | Top BLAST hits | BLAST %ID | Phylum;Class;Order |
| --- | --- | --- | --- | --- | --- | --- |
| OTU.6 | 3.62 | 7 | 3, 7, 14 | *Cellvibrio fulvus* | 100.0 | *Proteobacteria*<br>*Gammaproteobacteria*<br>*Pseudomonadales* |
| OTU.600 | 3.48 | 30 | 30 | No hits of at least 90% identity | 80.37 | *Planctomycetes* *Planctomycetacia*<br>*Planctomycetales* |
| OTU.6062 | 4.83 | 30 | 30 | *Dokdonella* sp. DC-3,<br>*Luteibacter rhizovicinus* | 97.26 | *Proteobacteria*<br>*Gammaproteobacteria*<br>*Xanthomonadales* |
| OTU.627 | 4.43 | 14 | 14 | *Verrucomicrobiaceae* bacterium DC2a-G7 | 100.0 | *Verrucomicrobia* *Verrucomicrobiae*<br>*Verrucomicrobiales* |
| OTU.633 | 3.84 | 30 | 30 | No hits of at least 90% identity | 89.5 | *Proteobacteria* *Deltaproteobacteria*<br>*Myxococcales* |
| OTU.638 | 4.0 | 30 | 30 | *Luteolibacter* sp. CCTCC AB 2010415,<br>*Luteolibacter algae* | 93.61 | *Verrucomicrobia* *Verrucomicrobiae*<br>*Verrucomicrobiales* |
| OTU.64 | 4.31 | 14 | 7, 14, 30 | No hits of at least 90% identity | 89.5 | *Chloroflexi* *Herpetosiphonales*<br>*Herpetosiphonaceae* |
| OTU.663 | 3.63 | 30 | 30 | *Pirellula staleyi* DSM 6068 | 90.87 | *Planctomycetes* *Planctomycetacia*<br>*Planctomycetales* |
| OTU.669 | 3.34 | 30 | 30 | *Ohtaekwangia koreensis* | 92.69 | *Bacteroidetes* *Cytophagia*<br>*Cytophagales* |
| OTU.670 | 2.87 | 30 | 30 | *Adhaeribacter aerophilus* | 91.78 | *Bacteroidetes* *Cytophagia*<br>*Cytophagales* |
| OTU.766 | 3.21 | 14 | 14, 30 | *Devosia insulae* | 99.54 | *Proteobacteria* *Alphaproteobacteria*<br>*Rhizobiales* |
| OTU.83 | 5.61 | 14 | 7, 14, 30 | *Luteolibacter* sp. CCTCC AB 2010415 | 97.72 | *Verrucomicrobia* *Verrucomicrobiae*<br>*Verrucomicrobiales* |
| OTU.862 | 5.87 | 14 | 14 | *Allokutzneria albata* | 100.0 | *Actinobacteria* *Pseudonocardiales*<br>*Pseudonocardiaceae* |
| OTU.899 | 2.28 | 30 | 30 | *Enhygromyxa salina* | 97.72 | *Proteobacteria* *Deltaproteobacteria*<br>*Myxococcales* |
| OTU.90 | 2.94 | 14 | 14, 30 | *Sphingopyxis panaciterrae*,<br>*Sphingopyxis chilensis*,<br>*Sphingopyxis* sp. BZ30,<br>*Sphingomonas* sp. | 100.0 | *Proteobacteria* *Alphaproteobacteria*<br>*Sphingomonadales* |
| OTU.900 | 4.87 | 14 | 14 | *Brevundimonas vesicularis*,<br>*Brevundimonas nasdae* | 100.0 | *Proteobacteria* *Alphaproteobacteria*<br>*Caulobacteriales* |
| OTU.971 | 3.68 | 30 | 30 | No hits of at least 90% identity | 78.57 | *Chloroflexi* *Anaerolineae*<br>*Anaerolineales* |
| OTU.98 | 3.68 | 14 | 7, 14, 30 | No hits of at least 90% identity | 88.18 | *Chloroflexi* *Herpetosiphonales*<br>*Herpetosiphonaceae* |
| OTU.982 | 4.47 | 14 | 14 | *Devosia neptuniae* | 100.0 | *Proteobacteria* *Alphaproteobacteria*<br>*Rhizobiales* |

<sup>a</sup> Maximum observed $\log_2$ of fold change.
<sup>b</sup> Day of maximum fold change.
<sup>c</sup> All response days.

